# MAIT cell enrichment in Lynch syndrome is associated with immune surveillance and colorectal cancer risk

**DOI:** 10.1101/2025.08.22.671764

**Authors:** Hairu Yang, Michaela Dungan, Keely Beyries, Xin Wang, Robert Kilpatrick, Baron Chen, Sangmi Oh, Marissa Berkowitz, David Smith, Sergei B Koralov, Jordan Axelrad, Christopher J. Lengner, Nicole Belle, Meenakshi Bewtra, Bryson W. Katona, Ken Cadwell

## Abstract

Tissue microenvironment characteristics associated with elevated risk of colorectal cancer (CRC) in Lynch syndrome (LS) are poorly characterized. We applied the multimodal single cell sequencing platform ExCITE-seq to define the colonic cellular composition and transcriptome of LS carriers with and without a history of CRC compared with general population controls. Our analysis revealed widespread remodeling in LS that included striking expansion of epithelial stem and progenitor cells, and loss of fibroblast populations. Although clonally expanded and terminally exhausted CD8 T cells were more prominent in individuals with a history of CRC, LS carriers without CRC displayed enrichment of cytotoxic mucosal-associated invariant T (MAIT) cells associated with *CCL20* expression in epithelial progenitors, validated by orthogonal techniques including demonstration of a protective function in a murine model of CRC. These findings highlight cellular features that distinguish LS carriers and suggest a protective role of MAIT cells in human CRC surveillance.

## INTRODUCTION

Colorectal cancer (CRC) is the third most common cancer and the second leading cause of cancer related death worldwide^1^. Lynch syndrome (LS) is the most common hereditary cause of CRC, affecting 1 in 279 individuals (approximately 1.2 million Americans) and conferring a lifetime CRC risk of up to 60%^2–7^. Similar to sporadic cancer, males with LS have a higher risk of CRC compared to females^8,9^. The increased risk of cancer in LS results from pathogenic germline variants (PGVs) in mismatch repair (MMR) genes *MLH1, MSH2, MSH6,* and *PMS2* or in *EPCAM,* which leads to methylation and subsequent silencing of *MSH2*^10^. These PGVs generally confer increased susceptibility to cancers by impairing the DNA mismatch repair system, a critical mechanism for correcting DNA errors generated during replication, leading to the accumulation of mutations that drive tumorigenesis^10–12^. Compared with the substantial progress in understanding how PGVs in MMR genes underlie cancer susceptibility in LS carriers, much less is known about how alterations in the tissue microenvironment contribute to CRC initiation and progression in this high-risk population^7,13^. A better understanding of the colorectal tissue properties in individuals with LS may provide insight into the factors distinguishing those who progress to CRC at a younger age.

The compositions and roles of immune cell types and their interactions with stromal and epithelial cells differ across CRC sub-types and stages^14–16^. In cases of sporadic CRC with high mutational burden, a robust CD8+ T cell response is generated against neoantigens followed by upregulation of exhaustion markers such as PD-1^16^. In contrast, adenomas caused by *adenomatous polyposis coli* (*APC*) PGVs display limited T cell infiltration due to interference from β-catenin signaling, and as the cancer progresses, regulatory T cells (Tregs) in the tumor microenvironment inhibit CD8+ T cells^17,18^. Although heighted immune activity is generally critical for anti-tumor responses, Th17-driven chronic inflammation can promote early tumorigenesis while also restricting further CRC progression^15,16,19^. Also, myeloid populations such as macrophages and neutrophils have been implicated in both tumor-promoting and tumor-suppressive roles, depending on their polarization and microenvironmental cues^19–21^. Endothelial cells, fibroblasts, and the epithelium participate by controlling access and activation of these immune infiltrates through establishing the structural environment, regulating hypoxia, and secreting chemokines and growth factors^14,15,22,23^. The design and success of cancer immunotherapy depends on these context-specific properties of the tissue microenvironment.

Our understanding of the immune landscape in LS-associated CRC remains limited, particularly during the early, pre-cancerous stages. A prior study using histology and bulk RNA sequencing revealed an enrichment of CD8+ immune cells in the unaffected colonic mucosa of cancer-free individuals with LS compared to those with CRC^24^. Follow up of LS carriers demonstrated a correlation between T cell infiltration of the tissue and time to subsequent tumor occurrence^24^, highlighting the importance of characterizing the immune microenvironment of the colon in this population. Given CD8 T cell heterogeneity, the implication of CD8 enrichment in the LS colon requires further elucidation. Also, the interaction of these T cells with other cells remains unexplored. To address this gap in knowledge, we performed multimodal single cell sequencing on colonic biopsies from LS carriers with or without a history of CRC. Compared with the general population, our analysis revealed that LS carriers exhibit extensive alterations in cellular compositions and transcriptional programs across the epithelial, stromal, and immune compartments. In LS carriers with a history of CRC, we observed pronounced TCR clonal expansion, while sex-specific differences were evident in B cell populations in both groups of LS carriers. Strikingly, mucosal-associated invariant T (MAIT) cells with a cytotoxic gene expression profile were enriched in the colons of LS carriers, especially in individuals without a cancer history. This enrichment was associated with expression of the chemokine *CCL20* by epithelial stem and progenitor cells. Functionally, increasing MAIT cells in a murine model of CRC reduced tumor burden and improved survival. Together, our findings reveal remodeling of the colonic tissue landscape in LS and implicate MAIT cells in CRC surveillance.

## RESULTS

### Expansion and transcriptional reprogramming of epithelial progenitors in the colons of LS carriers

Expanded Cellular Indexing of Transcriptomes and Epitopes by sequencing (ExCITE-seq) is a single-cell multiomic approach that integrates transcriptome and surface protein profiling at single-cell resolution, along with T and B cell receptor (TCR and BCR) sequencing for lymphocytes^25^. We performed ExCITE-seq on fresh, non-tumor proximal colonic biopsies collected from three age-and sex-matched groups: 10 LS carriers without CRC (LS), 6 LS carriers with a prior history of CRC (disease-free for over two years and not on treatment at the time of collection; LS-CRC), and 8 non-LS controls (undergoing routine screening colonoscopy; CON) (Figure 1A; Figure S1A-B; Table S1). Human CD4+ T cells were included during sequencing runs as an internal control for batch corrections^25,26^. After quality control, we recovered 30,501 total cells corresponding to an average of ∼1000 cells per individual (Figure S1C). Principal Component Analysis (PCA) of global gene expression profiles showed separation of CON from LS carriers (LS and LS-CRC) (Figure S1D). Through the integration of surface markers and transcriptome, we identified major cellular clusters corresponding to epithelial cells, stromal cells, T cells, B cells, and myeloid cells, each defined by distinct feature gene expression^27–29^ (Figure 1B, S1E). UMAP visualization demonstrated that major cell types were shared across samples without individual- or group-specific clustering, indicating successful batch integration and harmonization across datasets (Figure S1F-G). Notably, we observed a marked expansion of the epithelial compartment in LS carriers, especially in LS-CRC, compared with CON (Figure 1B, S1H).

**Figure 1.**
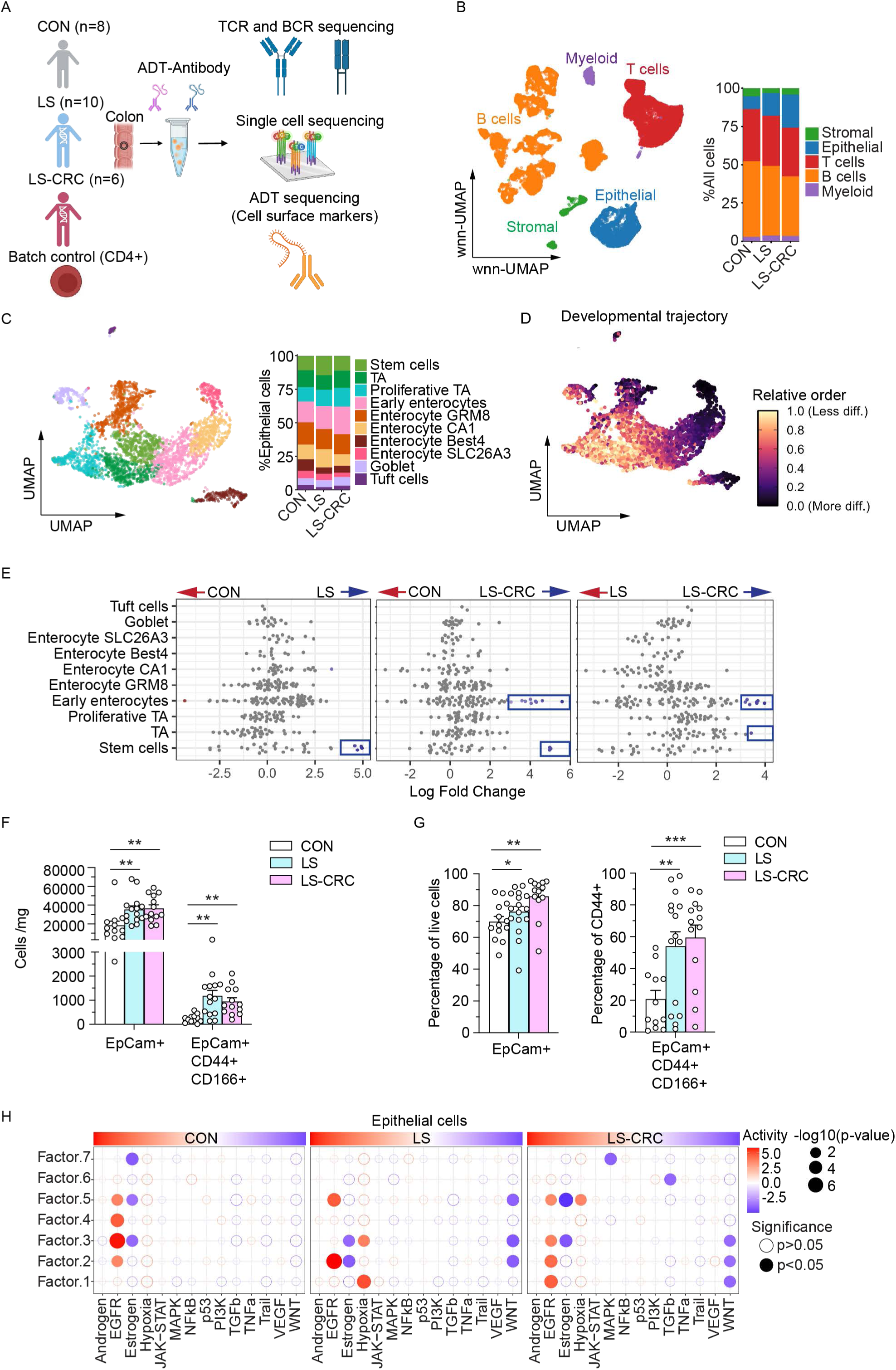
ExCITE-seq profiling of proximal colonic tissues reveals enrichment of epithelial progenitors in LS carriers. (A) Overview of the study design. Profiling of human colonic tissue from indicated subjects by ExCITE-seq, which simultaneously captures the surface epitope information using a cocktail of 60 oligo-conjugated antibody-derived tags (ADTs) and transcriptome of individual cells, along with the V(D)J repertoire of T cell and B cell antigen receptors of the same cell for lymphocytes. Isolated human CD4+ cells were included as batch control. Control (CON); Lynch syndrome without colorectal cancer (LS); Lynch syndrome with a prior colorectal cancer (LS-CRC). (B) wnn-UMAP embedding of cell surface markers and scRNA-seq from all cells (left panel) and bar graph indicating the percentage of each cell type in indicated individual groups (right panel). (C) UMAP showing the scRNA-seq from epithelium (left panel) and bar graph indicating the percentage of each cell type in indicated subject groups (right panel). (D) UMAP plot overlaid with CytoTRACE scores to show the developmental trajectory. (E) Beeswarm plot of adjusted log2 fold changes in abundance between experimental groups of cellular neighbourhoods (Nhoods) identified by Milo. Each dot represents a Nhood cluster and those displaying differential abundance (FDR < 0.1) are coloured in blue or red. (F) Absolute counts of EpCAM+ and EpCAM+CD44+CD166+ cells per mg of colorectal tissue, measured by flow cytometry. (G) Proportions of EpCAM+ and EpCAM+CD44+CD166+ cells as percentages of total live cells and EpCAM+CD44+ live cells, respectively. (H) Pathway enrichment in epithelial cells was analyzed using LIANA+ (Ligand-Receptor Inference Analysis). Factors in each row represent distinct groups of cell-cell communication and each dot represents a signaling pathway enriched within a specific factor. Dot color indicates pathway activity level, while size corresponds to statistical significance (−log₁₀ p-value, Holm correction). Dots represent one individual and bars show mean ± SEM for (F) and (G). Statistical significance for (F) and (G) was determined by one-way ANOVA (Fisher’s LSD post hoc test, *p < 0.05, **p < 0.001).

To characterize the expansion of the epithelial compartment, we performed sub-clustering analysis annotated by canonical epithelial lineage markers (Figure S2A) and identified three progenitor like clusters (stem cells, transit-amplifying (TA) cells, and proliferative TA), five clusters of absorptive enterocytes (Early enterocytes, Enterocyte GRM8, Enterocyte CA1, Enterocyte Best4, and Enterocyte SLC26A3), and two clusters of secretory epithelial cells (goblet cells and tuft cells) (Figure 1C, S2A). The UMAP plot overlaid with CytoTRACE scores^30^ revealed the developmental potential of epithelial clusters (Figure 1D). Stem/progenitor cells were positioned with higher CytoTRACE scores, reflecting a less differentiated state, while enterocytes at the periphery exhibited lower scores, indicating terminal differentiation (Figure 1D). Pseudotime analysis and fate probability mapping using CellRank^31^ demonstrated clear developmental trajectories from progenitor populations toward differentiated enterocyte subtypes (Figure S2B and S2C).

To investigate the cell compositional changes that may explain the epithelial expansion in LS and LS-CRC groups, we employed the computational strategy Milo for differential abundance testing^32^. Milo increases statistical rigor and avoids inherent biases that arise from cell-type clustering by grouping cells into overlapping neighborhoods based on similar gene expression. Using this approach, we identified stem cells to be enriched in LS and LS-CRC groups compared with CON (Figure 1E). The LS-CRC group was distinguished from LS by additional increases in TA cells and early enterocytes (Figure 1E). When considering the observation that LS carriers have a general expansion of all epithelial cells (Figure 1B), these findings suggest that LS and LS-CRC exhibit substantial epithelial remodeling characterized by increases in stem/progenitor cells. To validate these findings, we quantified total epithelial cells (EpCAM+) and epithelial stem/progenitor (EpCAM+CD44+CD166+) by flow cytometry in colorectal tissue samples from an expanded cohort that included additional age- and sex-matched subjects^33,34^ (Table S1). We observed two-fold or greater increases in EpCAM+ cells and EpCAM+CD44+CD166+ cells in LS carriers compared with CON (Figure 1F-G; S2D). Thus, epithelial and stem/progenitor populations are increased in the LS colon, even in histologically normal tissues.

To investigate the epithelial tissue microenvironment and its association with signaling pathways, we applied the LIANA+ (Ligand-Receptor Inference Analysis) framework to identify signaling pathways associated with cell-cell communication^35^. This analysis revealed enrichment of hypoxia pathway and downregulation of WNT pathway in LS and LS-CRC (Figure 1H). Differentially expressed gene (DEG) analysis revealed increases in transcripts associated with inflammation such as *IFITM1*, *LCN2*, and *REG1A* across multiple epithelial subtypes in LS carriers compared with CON and decreased expression of genes associated with epithelial homeostasis and differentiation, including AP-1 transcription factors (Figure S2E; Table S2). LS-CRC was further distinguished from LS by altered expression of factors associated with progenitor maintenance, developmental reprogramming, and tumorigenesis, including upregulation of *THRB*^36^ (Figure S2E; Table S2). Taken together, these findings indicate that composition and gene expression of the colonic epithelium is extensively altered in LS carriers.

### Lynch syndrome is associated with large-scale remodeling of the stromal compartment

Sub-clustering of the stromal compartment identified three fibroblast subtypes (cCF1 - colonic crypt fibroblasts 1, CTF - crypt top fibroblasts, and fibroblasts MYOCD), two endothelial subtypes (lymphatic endothelium and blood endothelial cells), pericytes, smooth muscle cells, and glial cells (Figure 2A, S3A). The relative abundance of these populations differed markedly across CON, LS, and LS-CRC (Figure 2A). Notably, cCF1 fibroblasts were substantially decreased in LS and LS-CRC based on differential abundance testing by Milo (Figure 2B). Located at the crypt base, cCF1s support epithelial stem cells by secreting WNT ligands and BMP inhibitors^37^. LIANA+ identified cCF1 as a major node for intercellular communication with epithelial populations, which was attenuated in LS carriers (Figure 2C). In contrast, blood endothelial cells were increased in both LS and LS-CRC (Figure 2B). DEG analysis identified a general downregulation of AP-1 transcription factors in fibroblast and endothelial populations (Figure 2D; Table S3). However, unlike the epithelium, inflammatory pathways including TNFα pathways, were downregulated in LS carriers (Figure S3B). When taken together, these results indicate that stromal reprogramming occurs in LS carriers.

**Figure 2.**
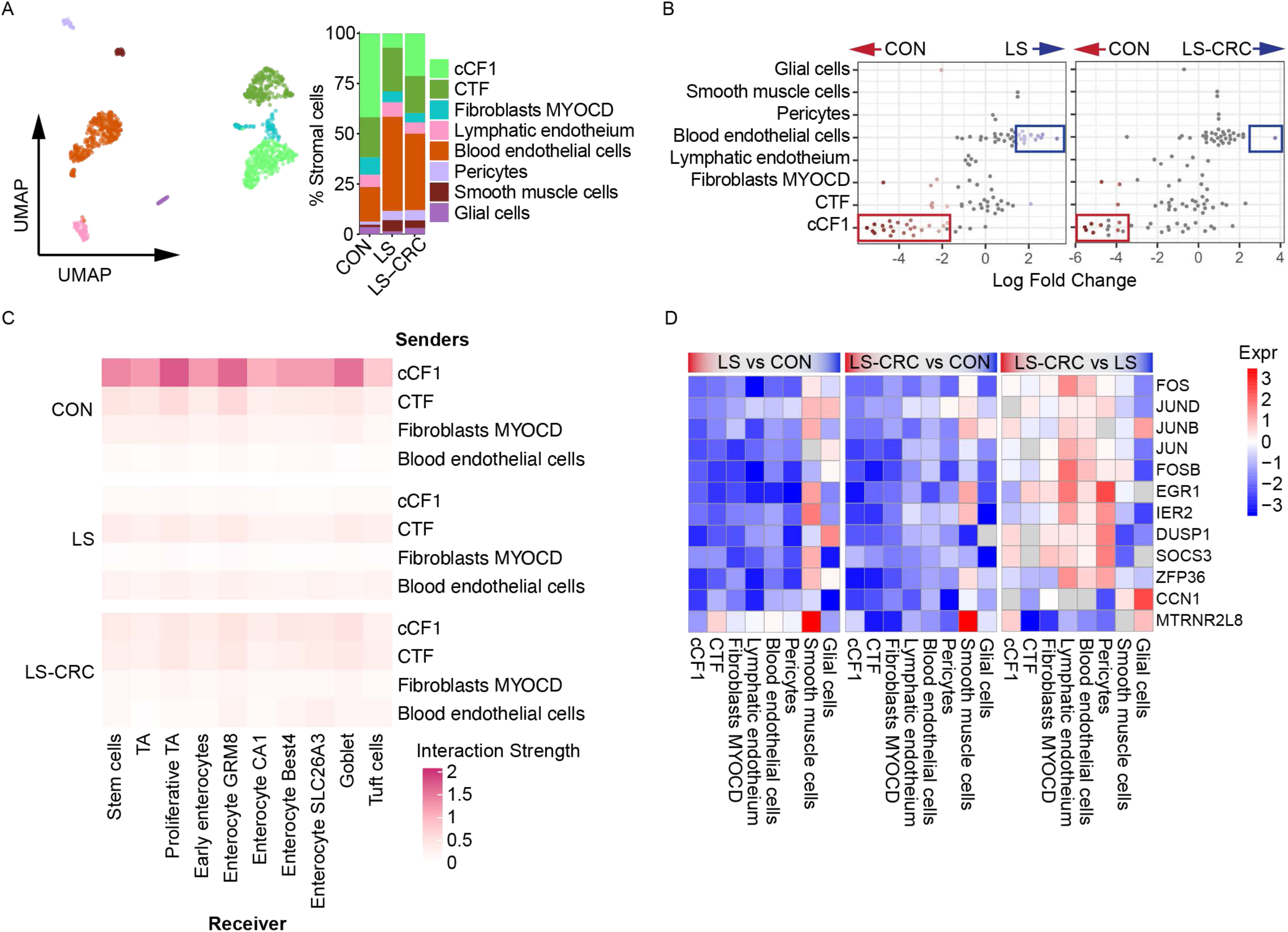
Lynch syndrome is associated with large-scale remodeling of the stromal compartment. (A) UMAP projection (left) and bar graph showing relative abundance (right) of stromal cell types in CON, LS, and LS-CRC. (B) Beeswarm plot of adjusted log2 fold change in abundance in CON vs LS and CON vs LS-CRC by Milo of cellular Nhoods identified by Milo. Differential abundance Nhoods (FDR < 0.1) are coloured in blue or red. (C) Heatmap depicting intercellular communication strength from stromal subtypes (senders) to epithelial cells (receivers), inferred by LIANA+. Interaction strength is normalized by sender cell abundance, and color intensity reflects the interaction strength. (D) Top DEGs per stromal subtype comparing LS vs CON, LS-CRC vs CON, and LS-CRC vs LS. Rows represent genes, and columns represent stromal subtypes. Color intensity denotes log₂ fold change (logFC) values, with upregulation in red and downregulation in blue. Non-significant genes (p-value > 0.05) are shaded in gray. Full statistical results including adjusted p-values (Benjamini-Hochberg correction) are provided in Table S3.

Although myeloid cells were relatively sparse, we identified two monocyte populations (Mono FTL, Mono CREM), two macrophage populations (macrophages C1QB, macrophages SLC8A1), and three dendritic cell subtypes (DC CD74, DC COTL1, DC FLT3) (Figure S3C-D). Differences in their relative abundances across groups were not significant. Therefore, we focused on lymphocytes for subsequent investigation.

### Sex differentially affects B cells in LS carriers

Sub-clustering of the B cell compartment identified six distinct subtypes: B naïve, B memory, and four plasma cell populations (IgG plasma cells (PC), IgA1 PC, IgA2 PC, IgA3 PC) (Figure 3A, S4A). Although the relative abundance of these B cell subtypes showed no significant differences across groups, we observed differential transcriptional patterns such as plasma cell-specific upregulation of long noncoding RNAs (lncRNAs) (*LINC00355*, *AC004556.3*, *AC105402.3*, and *LERFS*) in LS-CRC and *SFRP1* (Secreted Frizzled-Related Protein 1) in LS (Figure 3B; Table S4). In addition, we observed downregulation of immune response genes associated with antigen presentation, BCR signaling, FcγR-mediated pathways, and complement cascade in LS carriers, especially in LS-CRC (Figure 3B; Figure S4B). Unexpectedly, PCA revealed stratification by sex within the LS and LS-CRC groups but not in the CON (Figure 3C). Reanalysis of the relative abundance data for the B cell populations by sex revealed that the percentage of B memory cells were enriched and IgA2 plasma cells were reduced in females compared to males in LS and LS-CRC, which was confirmed by ratio of observed to expected cell numbers (Ro/e) analysis (Figure 3D; Figure S4C-D). When we compared gene expression differences between male and female subjects, the top differentially regulated genes were sex chromosome-linked such as *TTTY10*, *ZFY*, and *USP9Y* enriched in males, and *XIST* and *TSIX* enriched in females (Figure S4E; Table S5). GSEA revealed a potential reduction in the complement and activation signatures in LS females, which are normally seen enriched in CON group females (Figure 3E). Analyses of BCR clonal diversity, sharing, and occupancy showed comparable profiles across the three subject groups (Figure 3F, S5 A-C). Stratification of the BCR analyses by sex did not reveal major differences between males and females (Figure S5D-F). Thus, sex mainly impacts the abundance of B memory and IgA2 plasma cells in LS carriers and may have additional contributions to the transcriptomes of B cells.

**Figure 3.**
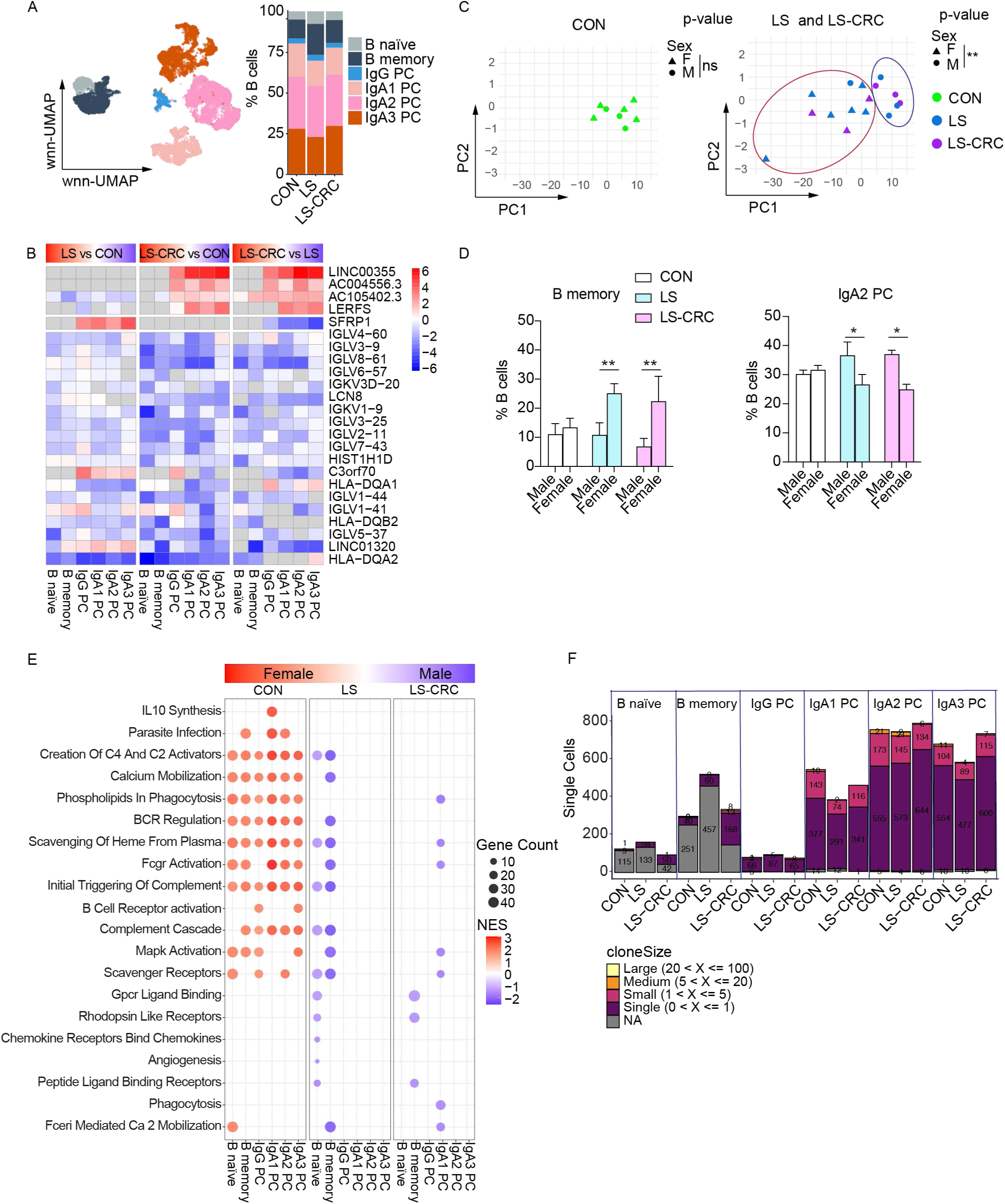
Sex differentially affects B cells in LS carriers. (A) wnn-UMAP projection (left) and bar graph showing relative abundance (right) of B cell subtypes in CON, LS, and LS-CRC. (B) Top DEGs per B cell subtype comparing LS vs CON, LS-CRC vs CON, and LS-CRC vs LS. Rows represent genes and columns represent B cell subtypes. Color intensity denotes logFC values, with upregulation in red and downregulation in blue. Non-significant genes (p-value > 0.05) are shaded in gray. Full statistical results including adjusted p-values (Benjamini-Hochberg correction) are provided in Table S4. (C) PCA plot based on transcriptional profiles of B cells. Each dot represents an individual. (D) Frequency of B memory cells and IgA2 plasma cells stratified by sex comparing CON, LS, and LS-CRC. Bars represent mean ± SEM. Statistical significance was determined by two-way ANOVA (Holm-Šidák post hoc test, *p < 0.05, **p < 0.001). (E) GSEA of B cells stratified by sex comparing CON, LS, and LS-CRC. Each dot represents an enriched pathway for a given B cell subtype, with color indicating the normalized enrichment score (NES) and dot size reflecting the number of genes contributing to the differential regulation (Gene count). Enrichment was performed using curated gene sets related to immune and signaling pathways. (F) Bar graph showing the number and size of clones in each B cell subtypes across groups after downsampling (2,477 cells per group) to normalize cell numbers across groups.

### T cell landscape and clonal dynamics are altered in LS carriers

By integrating cell surface staining markers and gene expression profiling, we identified distinct sub-clusters within the T cell super-cluster: CD4+ T cells (Naïve, Central memory, Th Progenitors, Th17 Progenitors, Th17 CCL20, Th17 Activated TNF, Th1, and Treg), CD8+ T cells (Stem central memory, Resident memory RA, Effector memory, Effector GZMK, Intermediate exhausted (Texh-int), Terminally exhausted (Texh-term), and MAIT), and innate lymphoid subtypes (ILC3s and tissue-resident NK (tNK) cells) (Figure 4A; Figure S6A). The distribution of these T cell clusters varied across groups (Figure 4A). Milo analysis largely confirmed the Ro/e data and highlighted enrichment of CD8 stem central memory and MAIT cells and a decrease in the CD4 Th17 cluster in LS compared to CON (Figure 4B). LS-CRC displayed an increase in CD8 Texh-int cells and a marginal elevation in MAIT cells (Figure 4B). The Th17 CCL20 cluster in LS and LS-CRC groups exhibited upregulation of genes associated with antigen presentation (*HLA-DQB1*, *HLA-DRB1*, *TRBV18*), inflammatory signaling (*IL22*, *GPR25*, *ALPK2*, *HDGFL3*, *ARL17B*), transcriptional control and differentiation (*SOX5*, *LINC02384*, *MIR181A1HG*), and cellular stress protection (*HSPA1A*, *HSPA1B*) compared with CON (Figure S6B; Table S6). Similarly, pathway analysis by GSEA revealed that Th17 clusters in LS carriers upregulated cell adhesion, antigen presentation, cytokine signaling, and stress protection (Figure S6C). In contrast, CD8 effector cells exhibited downregulation of multiple key genes involved in tumor-suppressive transcriptional regulation (*HIST1H1C*, *ZNF331*, *KLF6*, and *BTG1*) and immune activation and trafficking (*CXCR4*, *CD69*, and *TNFAIP3*) in LS carriers, which was more severe in LS-CRC (Figure S6D; Table S6). Supporting these transcriptional changes indicative of reduced function, GSEA revealed suppression of pathways critical for CD8 T cell activation and effector function, including FGFR signaling, Wnt/β-catenin signaling, and the antimicrobial peptides pathway, especially in LS-CRC (Figure S6E).

**Figure 4.**
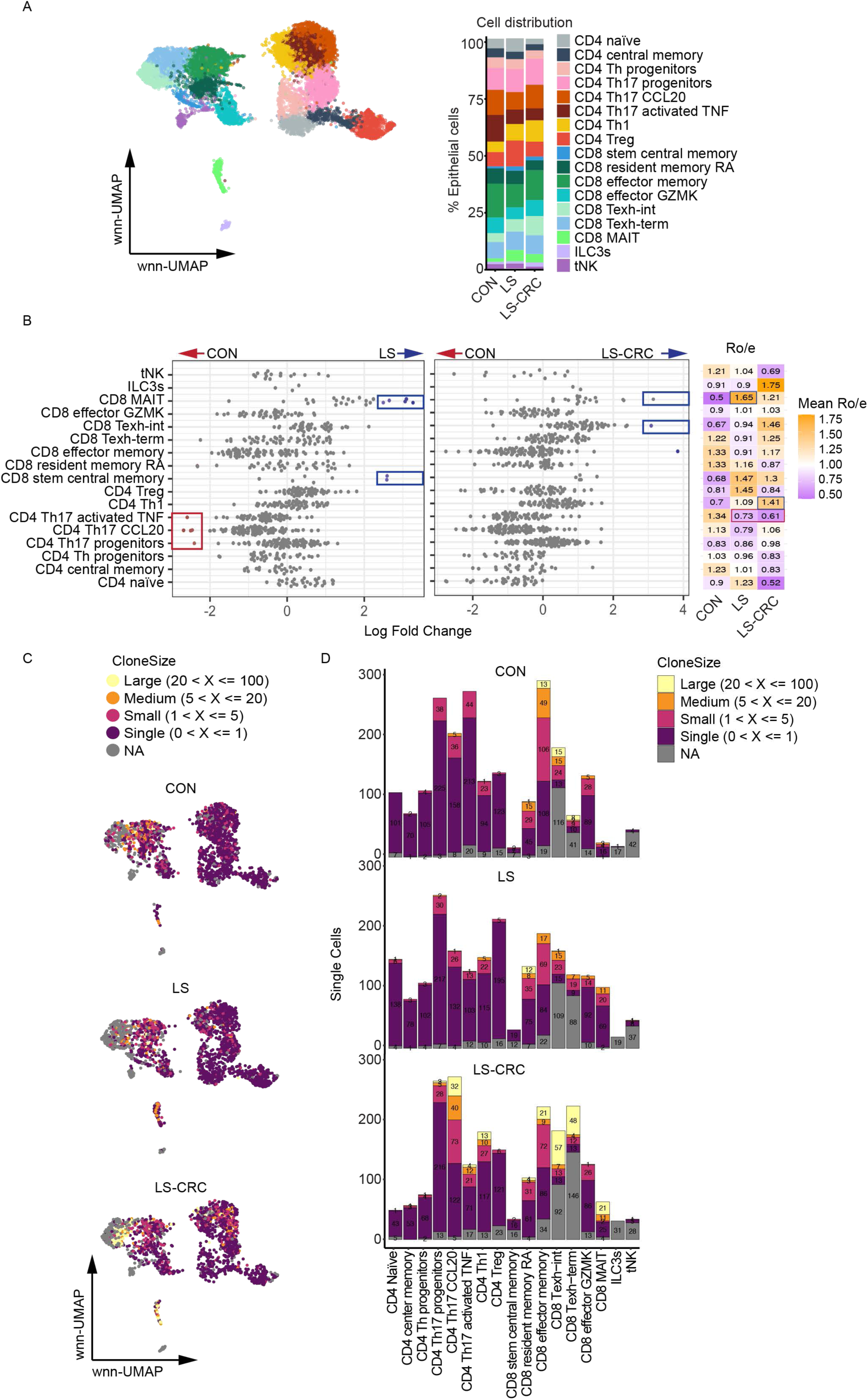
T cell landscape and clonal dynamics are altered in LS carriers. (A) wnn-UMAP projection integrating transcriptomic and surface protein data (left) and bar graph showing relative abundance (right) of T cell subtypes in CON, LS, and LS-CRC. (B) Beeswarm plot of adjusted log2 fold change in Nhood abundance comparing CON vs LS and CON vs LS-CRC by Milo. Each dot represents a Nood cluster and those displaying differential abundance (FDR < 0.1) are colored in blue or red. Heatmap shows ratio of observed to expected cell number (Ro/ e) scores for T cell subtypes corresponding to those in the same row as the beeswarm plot comparing CON, LS, and LS-CRC. Values >1 indicate overrepresentation relative to expected frequencies. Scores are color-scaled by mean Ro/e per group. Statistical significance of deviations was assessed using the Wilcoxon rank-sum test. Comparisons with p < 0.05 are color-coded in blue or red. (C) TCR clone size mapped onto wnn-UMAP plots of T cells from the indicated groups after downsampling (n = 2,191 cells per group). Colors represent clonal sizes: Large (yellow; 20 < X ≤ 100), Medium (orange; 5 < X ≤ 20), Small (purple; 1 < X ≤ 5), Single (dark red; 0 < X ≤ 1), and NA (gray). (D) Bar graphs displaying the distribution of TCR clone sizes across T cell subtypes in the indicated groups. The numbers inside the bars indicate the total cell counts for each subtype. Colors represent TCR clone sizes: Large (yellow; 20 < X ≤ 100), Medium (orange; 5 < X ≤ 20), Small (purple; 1 < X ≤ 5), Single (dark red; 0 < X ≤ 1), and NA (gray).

Examination of TCR repertoires showed that T cells in LS-CRC exhibit more clonal expansion relative to CON or LS (Figure S7A-B). Mapping the clone sizes onto the T cell UMAP clusters indicated that these expanded clones represented Th17 CCL20, Th17 Activated TNF, and Th1 clusters among CD4 subtypes and Effector memory, Texh-int, Texh-term, and MAIT cells among CD8 subtypes (Figure 4C, D). Exhausted CD8 T cell clusters, Texh-int and Texh-term shared overlapping TCR clones, which was especially pronounced in LS-CRC (Figure S7C), potentially reflective of a chronic antigen-driven transition into different exhausted states. A substantial degree of clonal sharing was also observed between CD4 Th17 CCL20 and CD4 Th1 subtypes specifically in LS-CRC (Figure S7C), suggesting functional plasticity or transition between Th17 and Th1 lineages. A proportion of T cells in LS and LS-CRC groups carried complementarity-determining region 3 (CDR3) sequences known to recognize cancer-related neoantigens, suggesting ongoing immune surveillance even in non-tumor colonic tissue (Figure S7D).

### MAIT cells with cytotoxic potential are enriched in LS carriers without a history of CRC

MAIT cells were among the most statistically robust lymphocytes that were enriched in LS tissue and have not been sufficiently examined in the context of human CRC. Although MAIT cells were discovered over 25 years ago, they have only recently attracted attention, mostly in the context of infections where they promote host defense by recognizing microbial metabolites derived from riboflavin synthesis (e.g., 5-OP-RU)^38^. MAIT cells harbor semi-invariant TCRs recognizing microbially-derived products and self-ligands presented by the non-classical MHC-I molecule MR1^39^. In humans, MAIT cells exhibit tissue-specific variation in both abundance and functional capacity, reflecting their adaptation to distinct microenvironmental cues^40^. Whether MAIT cells are pro- or anti-cancer is unclear, and new self-ligands such as nucleobase adducts and sulfated bile acid are beginning to be identified^38,41–49^.

Quantification of MAIT cells by flow cytometry in additional subjects using anti-TRAV1-2 antibody and a 5-OP-RU MR1 tetramer^40,44,45^ (Figure 5A; Figure S8A; Table S1) confirmed that these cells were increased in LS carriers compared to CON (Figure 5B). The cell surface markers allowed us to identify three subpopulations of MAIT cells, which displayed similar proportions across all groups: TRAV1-2+ (around 60%), 5-OP-RU tetramer+ MAIT cells (around 35%), and TRAV1-2+/5-OP-RU+ double-positive cells (about 5%) (Figure 5A and 5C). The proportion of TRAV1-2 positive versus negative MAIT cells in our ExCITE-seq data was similar to this flow cytometry analyses (Figure 5D; Table S7). To gain insight into the functional capacity of MAIT cell subtypes, we analyzed the expression of key effector genes, including *GZMA*, *GZMK*, *PRF1* (*perforin*), *FASLG*, *TNFSF10*, along with the activation marker *CD69*^40^ (Figure 5E). The TRAV1-2+ MAIT cells displayed high expression of these markers (Figure 5E). When comparing these effectors across subject groups, we found that they were decreased in expression in LS-CRC MAIT cells compared with CON and LS (Figure 5F). Therefore, both LS and LS-CRC have increased numbers of MAIT cells, but those in LS-CRC may be less cytotoxic and/or functional.

**Figure 5.**
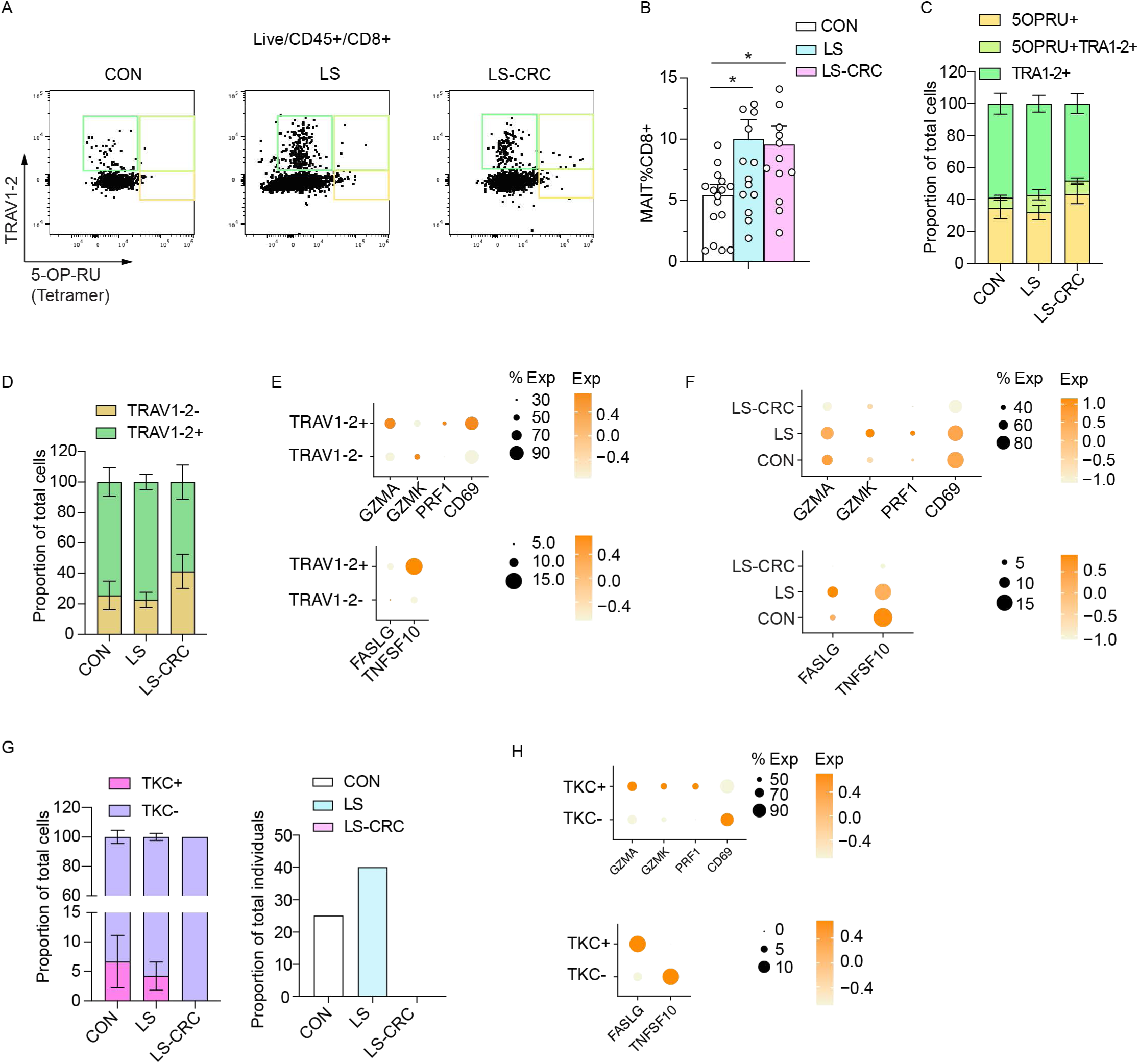
MAIT cells with cytotoxic potential are enriched in LS carriers without a history of CRC. (A) Representative flow cytometry plots of CD8+ MAIT cells identified by 5-OP-RU-loaded MR1 tetramers and anti-TRAV1-2 staining in colonic tissue from CON, LS, and LS-CRC. (B) Quantification of MAIT cells as a percentage of CD8+ T cells from (A). Statistical significance was determined by one-way ANOVA (Fisher’s LSD post hoc test, *p < 0.05, **p < 0.001). (C) Bar graphs showing the proportion of MAIT cells stained by 5-OPRU MR1 tetramers and anti-TRAV1-2 for CON, LS, and LS-CRC. (D) Proportion of TRAV1-2+ or TRAV1-2-MAIT cells based on ExCITE-seq analysis. (E) Dot plots displaying gene expression of cytotoxic markers (GZMA, PRF1, CD69) and apoptosis-related genes (FASLG, TNFSF10) in TRAV1-2+ versus TRAV1-2-MAIT cells from the ExCITE-seq dataset. (F) Comparison of MAIT cell cytotoxic and apoptosis-related gene expression in MAIT cells across CON, LS, and LS-CRC from the ExCITE-seq dataset. (G) Frequencies of TCR-defined killer cell (TKC) clones (TRBV12-4, TRBV24, TRBV25-1, TRBV28, and TRBV29-1) across three groups (Left panel). Proportion of individuals in each group carrying the TKC clones (Right panel). (H) Comparison of cytotoxic and apoptotic gene expression between TKC+ and TKC-subsets from the ExCITE-seq dataset.

MAIT cell carrying TRBV12-4, TRBV24, TRBV25-1, TRBV28, and TRBV29-1 rearrangements have been previously identified as having anti-tumor capabilities and are collectively referred to as tumor-killing clones (TKCs)^39,50^. We observed that these TKCs were present in both the CON and LS groups but notably absent in the LS-CRC group (Figure 5G). Furthermore, we found that the effector molecules, including *GZMA*, *GZMK*, *PRF1*, and *FASLG*, were upregulated in TKCs compared with other MAIT cell populations (Figure 5H; Table S8). Therefore, LS displays an increased presence of TKCs with a cytotoxic gene expression profile compared with LS-CRC.

### *CCL20+* stem and progenitor epithelial cells are enriched in LS carriers

To explore the interaction dynamics between epithelial cells and MAIT cells, we performed cell-to-cell communication analysis with a focus on chemokine signaling (Figure 6A). This analysis revealed that epithelial progenitors (Stem cells, TA, Proliferative TA, Early enterocytes, and Enterocyte GRM8) displayed markedly stronger signaling interactions with MAIT cells in the LS group compared to CON and LS-CRC (Figure 6A).

**Figure 6.**
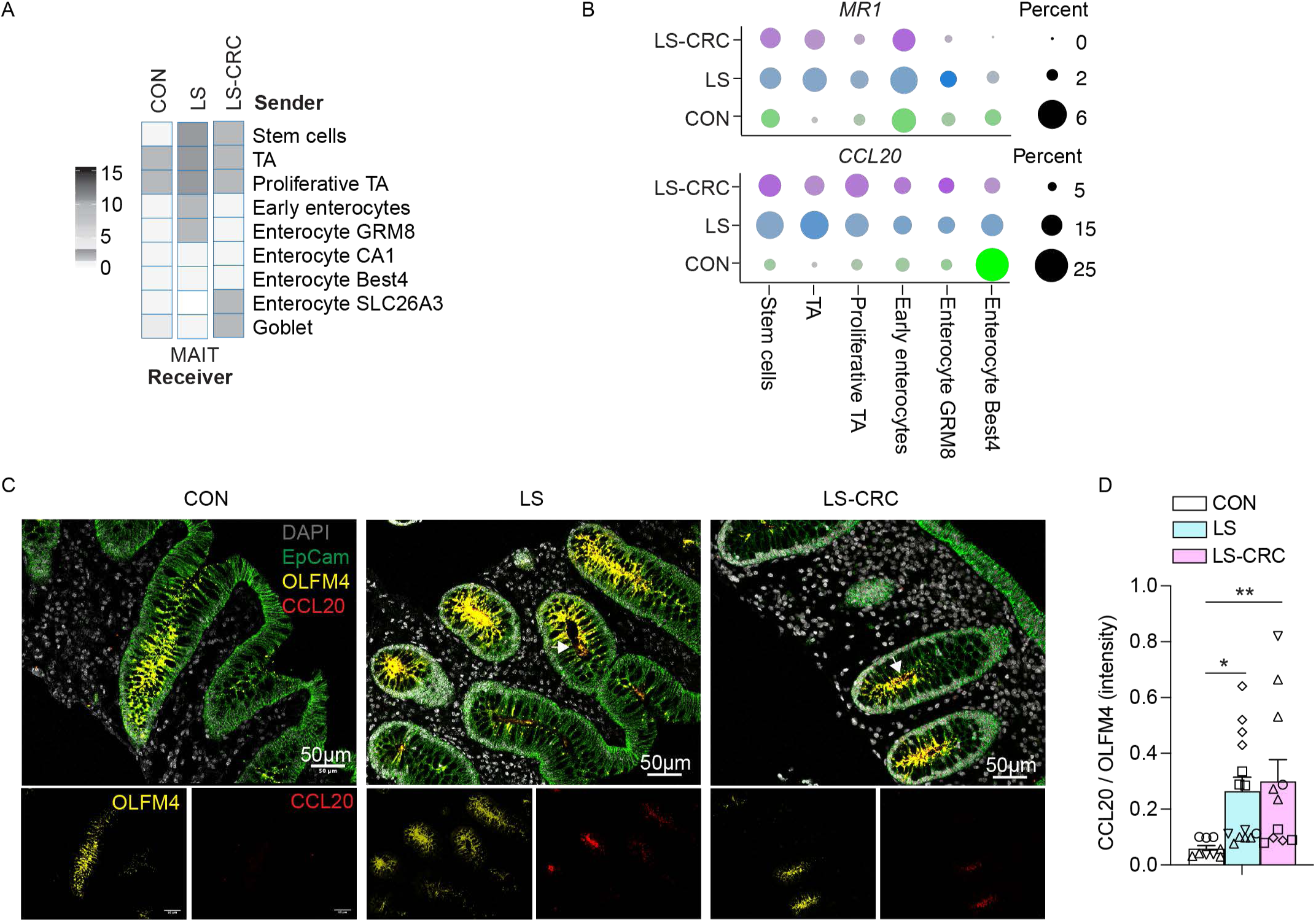
CCL20+ stem and progenitor epithelial cells are enriched in LS carriers. (A) Heatmap depicting predicted intercellular communication strength from epithelial subtypes (senders) to MAIT cells (receivers), inferred using the LIANA framework. Interaction intensity is represented by color scale. (B) Dot plots showing expression levels of MR1 (top) and CCL20 (bottom) across epithelial subtypes in each group. Dot size indicates the percentage of expressing cells; color intensity reflects expression level. (C) Representative immunofluorescence images of colonic crypts stained for CCL20 (red), OLFM4 (yellow), EpCAM (green), and DAPI (gray). CCL20 is enriched in OLFM4+ epithelial stem and progenitor cells in LS and LS-CRC tissues. (D) Quantification of the CCL20 to OLFM4 fluorescence intensity ratio. CCL20 expression is significantly increased in LS and LS-CRC compared to CON. Bars represent mean ± SEM. Identical symbols denote different area from same individuals (N 3). Statistical significance was determined by one-way ANOVA (Fisher’s LSD post hoc test, *p < 0.05, **p < 0.001).

Examination of known chemokines associated with MAIT cell trafficking revealed *CCL20* was specifically upregulated in in both LS and LS-CRC groups compared to the CON^38,51–53^ (Figure 6B; Figure S9A). Additionally, we observed increased expression of *MR1* within the same stem and progenitor populations (Figure 6B; Figure S9A). Consistent with these findings, UMAP projections and quantitative analysis demonstrated that epithelial cells positive for OLFM4, a stem and progenitor marker that is also associated with CRC^54–57^, co-expressed *CCL20*, which was more pronounced in LS and LS-CRC compared to CON (Figure S9B-C). To validate the observed expression patterns, we performed immunofluorescence staining of colonic tissue for CCL20, OLFM4 (epithelial progenitors), and EpCAM (epithelial cells). We observed robust CCL20 staining that was predominantly localized to OLFM4+ cells within the EpCAM+ epithelial compartment in LS and LS-CRC compared to CON (Figure 6C-D). Therefore, OLFM4+ epithelial cells exhibit increased CCL20 expression in the colons of LS carriers.

### Boosting MAIT cells is protective in a mouse model of CRC

A prior study demonstrated that MR1-deficiency increases the number of large tumors in a model of CRC in which mice are treated with azoxymethane (AOM) and dextran sodium sulfate (DSS)^44^. In contrast, we sought to address whether increasing MAIT cells above baseline, as observed in LS carriers, could confer protection. Intraperitoneal administration of 5-OP-RU in combination with CpG as an adjuvant was previously shown to expand MAIT cells^42^. Therefore, we repetitively administered 5-OP-RU with or without CpG to mice receiving a course of AOM and DSS (Figure 7A). Mice treated with 5-OP-RU, 5-OP-RU + CpG, or vehicle control experienced similar body weight loss after DSS exposure (Figure S10A). However, mice treated with 5-OP-RU or 5-OP-RU + CpG displayed improved survival compared with the vehicle control group (Figure 7B). Additionally, examination of the colons from the surviving mice indicated that treatment with 5-OP-RU or 5-OP-RU + CpG reduced tumor volumes (Figure 7C). Flow cytometry analysis showed that 5-OP-RU or 5-OP-RU + CpG treated mice displayed a substantial increase in MAIT cells in the tumor-free regions of the colon compared to vehicle controls (Figure 7D). In contrast, MAIT cells in the tumor tissue were comparable across three groups (Figure 7D). The percentage of MAIT cells in the tumor-free regions of the colon, but not tumor tissues, displayed an inverse correlation with tumor volume (Figure 7E). Together, these findings indicate that an increased presence of MAIT cells in the colon is protective against colorectal tumor formation.

**Figure 7.**
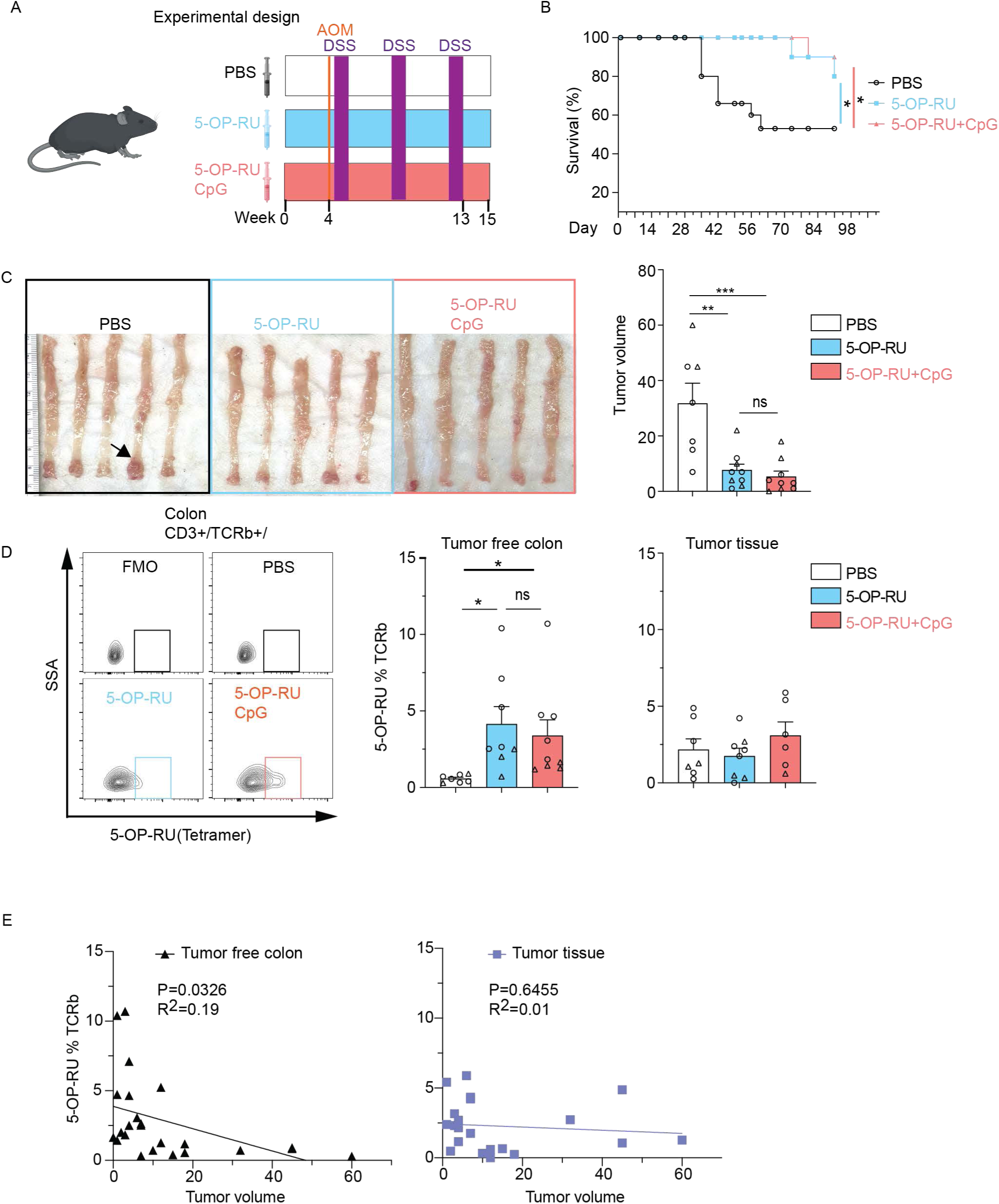
Increasing MAIT cells reduces colorectal tumor burden in AOM/DSS treated mice. (A) Experimental design: mice received twice weekly treatments with PBS (vehicle), 5-OP-RU, or 5-OP-RU + CpG adjuvant, followed by administration of AOM and DSS. (B) Survival curves of mice from (A) (n 10 per group). Log-rank (Mantel-Cox) test was used for statistical comparison. (C) Representative images of colons and quantification of tumors at endpoint in surviving mice. Arrow indicate visible tumors. (D) Flow cytometry analysis of MAIT cells (CD3+TCRβ+5-OP-RU tetramer+) in tumor free colon and colon tumor tissues. Fluorescence minus one (FMO) and PBS are shown as gating controls. (E) Scatter plots showing the relationship between MAIT cell frequency and tumor volume in tumor-free colonic tissue (left) and colon tumor tissue (right). Dots represent one individual mouse and bars show mean ± SEM for (C) and (D). Different symbols (triangles or circles) denote animals from independent experimental replicates. Statistical significance was determined by one-way ANOVA (Fisher’s LSD post hoc test, *p < 0.05, **p < 0.001, ns = not significant).

## DISCUSSION

Our comprehensive ExCITE-seq profiling with validation by flow cytometry and immunofluorescence microscopy uncovered extensive alterations in the proportions and functional state of multiple cell populations in the colonic tissue from LS carriers compared with the general population controls. Among notable differences were a substantial expansion in LS carriers of epithelial stem and progenitor cells skewed towards a pro-inflammatory gene expression profile that included expression of *CCL20*. Although enrichment of MAIT cells was also a general feature of LS carriers, MAIT cells in LS carriers without CRC were distinguished by their cytotoxic gene expression profile and TCR sequences previously associated with tumor killing. Consistent with these human data, inducing expansion of MAIT cells reduced colorectal tumor burden and improved survival in mice. These findings support a model wherein the colons of individuals with LS display a pro-inflammatory state at baseline, which includes expansion of CCL20-producing epithelial cells to mobilize MAIT cells that participate in anti-tumor surveillance. The degree to which cytotoxic activity is retained by these MAIT cells and other CD8 T cell populations recruited to the affected tissue is likely an important determinant of when individuals with LS develop CRC.

Several observations increase our confidence in the dataset we generated. First, the increase in the blood vessel endothelium and clonal expansion of exhausted CD8 T cells align well with the literature showing angiogenesis and lymphocyte enrichment in the colons of individuals with LS and/or CRC^24,58,59^. Also, epitope prediction analysis indicated that a subset of the TCRs recognize neoantigens in LS carriers, suggesting ongoing immune surveillance. LS-CRC individuals exhibited pronounced TCR clonal expansion of exhausted CD8+ cells and Th17 subtypes, features absent in those without CRC, aligning with prior reports showing pronounced clonal expansion within exhausted CD8+ and Th17 subsets^60–64^. Therefore, we believe the ExCITE-seq data will serve as a useful resource for the field.

CRC development is primarily driven by disruptions in core signaling pathways in epithelial cells, notably the aberrant activation of WNT and EGFR and the concurrent suppression of the TGFβ-BMP axis^14,65^. Our data revealed downregulation of WNT and TGFβ-BMP pathways in the epithelium. This profile coincided with depletion of cCF1 fibroblasts previously reported to be a source of WNT family ligands that contributes to stem and progenitor cell maintenance^37,66^. This potential disruption of the stem cell niche aligns with recent organoid models describing a transition from regenerative-like stem cells (revCSCs) to hyper-proliferative colonic stem states (proCSCs), driven by reduced WNT and TGFβ signaling^67,68^. We speculate that this revCSC-to-proCSC trajectory is occurring in the non-tumor colon in individuals with LS. Supporting this concept, stromal changes, specifically fibroblast loss and endothelial expansion, reflect a tissue microenvironment undergoing pre-malignant remodeling and immune cell infiltration^69–72^. An important future direction would be to determine how PGVs in MMR genes contribute to these disruptions in cellular networks.

Altogether, our study reveals that LS carriers exhibit coordinated remodeling of the epithelial, stromal, and immune compartments, even in histologically normal tissues. Notably, we provide evidence of a role for MAIT cells in human CRC. Given the elevated CRC risk in LS carriers and differences in CRC penetrance, risk stratification is important to guide colorectal surveillance strategies. Our findings reveal several markers, including MAIT cells and epithelial progenitors, that may facilitate improved risk stratification, guide surveillance strategies, and inform the development of intervention strategies in Lynch syndrome.

### Limitation of our study

ExCITE-seq represents a high-resolution platform but is difficult to scale. Therefore, many of the findings will require confirmation in larger cohorts. Additionally, longitudinal sampling will be required to directly associate MAIT cell abundance with clinical outcomes and determine whether *CCL20* expression by epithelial stem and progenitor cells precedes their enrichment. Also, MAIT cells are heterogenous; subsets and closely related unconventional T cell subsets such as MR1 like cells (also referred to as MR1-restricted T cells) are often difficult to distinguish. Therefore, although our analysis includes 5-OP-RU MR1 tetramers and bona fide MAIT cells, it remains possible that additional subsets of unconventional T cells are altered in the LS colonic tissue. Finally, although AOM/DSS treatment of mice is widely used to investigate CRC and provided evidence that increased presence of MAIT cells is protective, this model does not capture the genetic landscape of LS. Improvements in existing murine models of CRC-associated with LS will be invaluable for investigating the mechanism by which MAIT cells and other colonic tissue properties contribute to tumor surveillance.

## SUPPLEMENTAL INFORMATION

**Document S1. Figures S1–S10**

**Table S1. Human subjects**

**Table S2. DEGs in epithelial cells**

**Table S3. DEGs in stromal cells**

**Table S4. DEGs in B cells by cancer status**

**Table S5. DEGs in B cells by sex**

**Table S6. DEGs in T cells**

**Table S7. DEGs in TRAV1-2+ versus TRAV1-2-cells**

**Table S8. DEGs in TKC+ versus TKC-cells**

**Table S9. Key resources table**

## METHOD

### Patient enrollment and tissue sample collection

Human tissue samples were obtained through the LIMBO (Lynch Syndrome Mucosal Immune and MicroBiOme) study under an IRB-approved protocol (IRB#835032, University of Pennsylvania). Informed consent was obtained from all participants prior to enrollment, and all procedures were conducted in accordance with institutional and federal guidelines. The study included three cohorts: (1) Lynch syndrome (LS) carriers without a history of colorectal cancer (CRC), (2) Lynch syndrome carriers with a prior history of CRC (LS-CRC), and (3) age- and sex-matched controls without a diagnosis of LS. All LS and LS-CRC participants carried clinically confirmed pathogenic germline variants in a mismatch repair (MMR) gene (*MLH1*, *MSH2*, *MSH6*, or *PMS2*) or *EPCAM*. For LS-CRC participants, we selected individuals who had completed colorectal cancer (CRC) treatment at least two years prior to enrollment, were confirmed to be cancer-free, and were not receiving any ongoing cancer therapy at the time of sample collection. Control participants were recruited during routine screening colonoscopies and had no personal or family history of colorectal cancer, inflammatory bowel disease (IBD), autoimmune disorders, infectious colitis, or hereditary cancer syndromes. During colonoscopy, mucosal biopsies were obtained endoscopically using jumbo biopsy forceps of normal appearing colonic mucosa. Two to four biopsies were collected from both the proximal most colon and distal colon, immediately placed into cryovials containing chilled phosphate-buffered saline (PBS, ThermoFisher, Cat# 10010049), and transported on wet ice for prompt processing. To preserve native transcriptional states, tissue samples were cryopreserved within 2-4 hours using a freezing medium composed of 90% fetal bovine serum (FBS, PEAK, Cat# PS-FB2) and 10% dimethyl sulfoxide (DMSO, Sigma-Aldrich, Cat# D2650). Only histologically normal, non-neoplastic mucosal samples were retained for downstream single-cell analysis. Post hoc histopathological evaluation confirmed epithelial integrity and absence of active disease or inflammation.

### Single cell dissociation for ExCITE seq and flow cytometry

Frozen mucosal biopsies were rapidly thawed in a 37°C water bath for 2 minutes and immediately transferred to ice. Tissue was washed three times in 10 mL cold Hank’s Balanced Salt Solution (HBSS, Fisher Scientific, Cat# 14-175-103) by inversion. Samples were incubated in a 5 mL dissociation buffer containing PBS supplemented with 2.5 mM EDTA (Corning, MT-46034CI) and 1 mM DTT (Sigma, Cat# D0632-5G) for 90 minutes at 4°C on a tube rocker. Following incubation, tissue was washed with DMEM (ThermoFisher, Cat# MT-10-013-CV) containing 10 µM Y27632 (ROCK inhibitor, Sigma, Cat# Y0503-5MG) and vigorously shaken 100 times to release crypts. This shaking step was repeated 2-4 times until supernatants became visibly clear. All supernatants were pooled and centrifuged at 400g for 5 minutes at 4°C. After adding the 5 mL digestion solution composed of RPMI 1640 (ThermoFisher, Cat# 61870127) with 10% FBS, 1% Penicillin/Streptomycin (Sigma, Cat# P4333-100ML), 10 mM HEPES (Corning, Cat# 25-060-CI), Collagenase Type I (1 mg/mL, Sigma, Cat# C2139-1G), and DNase I (20 U/mL, Sigma, Cat# DN25-1G), the samples were shacked with speed 200RPM/Minute at 37°C for 30 minutes. Enzymatic digestion was quenched with 15 mL cold cRPMI and filtered through a 100 µm strainer (Corning, Cat#352360). Cell suspension was centrifuged at 400g for 5 minutes and resuspended in FACS buffer for downstream analysis. For ExCITE-seq, the live cells were further enriched by Dead Cell Removal Kit (Miltenyi Biotec, Cat#130-090-101) to ensure viability of all samples is at least 90%.

### ExCITE-seq

Single-cell suspensions (∼200,000 cells per sample) were resuspended in staining buffer composed of PBS with 2% bovine serum albumin (BSA, Sigma, Cat# P7030). Samples were centrifuged at 300–350 × g for 5 minutes at 4 °C, and the supernatant was carefully discarded. Cell pellets were then resuspended in 12.5 μL staining buffer supplemented with Fc receptor blocking reagent TruStain FcX™ (BioLegend, Cat# 422302) and incubated on ice for 10 minutes. Following Fc blocking, 12.5 μL of a 2X barcoded antibody cocktail was added directly to each tube. This cocktail contained a panel of oligonucleotide-conjugated antibodies for CITE-seq and a unique hashing antibody for each sample (Table S9). Cells were gently mixed and incubated on ice for 30 minutes. Post-incubation, cells were washed three times in staining buffer and a final time in staining buffer lacking BSA. Cells were filtered through a 70 μm strainer (Fisher Scientific, Cat# 22-363-548) to remove aggregates, then counted. Equal numbers of cells from each sample were pooled, centrifuged, and resuspended in PBS for subsequent loading onto the 10x Genomics Chromium Controller^25,26^.

### Single-cell multiomic data processing and analysis

Raw sequencing data were processed using the Cell Ranger software suite (10x Genomics) for sample demultiplexing, alignment to the human reference genome (GRCh38, Ensembl: http://useast.ensembl.org/Homo_sapiens), and quantification of unique molecular identifiers (UMIs). Feature barcoding count matrices, including antibody-derived tags (ADTs) and hashtag oligonucleotides (HTOs), were generated using the 10x Genomics Cloud Analysis service. Sample demultiplexing based on HTOs was performed using the HTODemux function implemented in Seurat v5.0.1. Quality control was performed to exclude low-quality cells and technical artifacts. Epithelial cells with fewer than 200 detected genes or more than 20% mitochondrial gene content were removed, while a 15% mitochondrial gene content threshold was applied to non-epithelial cell types. Doublets were identified and excluded using the scDblFinder package^73^. RNA expression data were normalized and batch-corrected using the Harmony package^74^.

### Differential gene expression analysis and pathway enrichment analysis

Differential gene expression analysis was performed using the FindMarkers function in Seurat (v5.0.1)^75^. For each comparison, normalized and batch-corrected RNA expression matrices were used. The Wilcoxon rank-sum test was applied with default parameters to identify differentially expressed genes between groups or clusters. Genes were considered significant if they met an adjusted p-value threshold of < 0.05 and an average log-fold change > 0.25, unless otherwise specified. P-values were adjusted using the Benjamini-Hochberg false discovery rate (FDR) correction. Pathway activity was assessed using the clusterProfiler R package with curated gene sets from MSigDB, including the Hallmark (H) and Reactome collections^76^. Enrichment analyses were conducted using gene set enrichment analysis (GSEA), and normalized enrichment scores (NES) were subsequently correlated with transcriptional signatures across conditions or cell types.

### TCR and BCR analysis

For B cell receptor (BCR) and T cell receptor (TCR) analysis, the scRepertoire R package^77^ (v2.0.3) was used to analyze V(D)J sequencing data processed via Cell Ranger VDJ (10x Genomics). Annotated contig files were imported and integrated with the single-cell Seurat object using the combineExpression() function. Clonotypes were defined based on identical Complementarity-Determining Region 3 (CDR3) sequences in both T (TRA and TRB) and B cells (IGH and IGK/IGL). Clonotype expansion, diversity metrics, TCR/BCR repertoire sharing, and clonal overlap were assessed across conditions (CON, LS, and LS-CRC) using scRepertoire, following downsampling to normalize cell numbers across groups. To assess potential antigen specificity, TCR sequence convergence analysis was performed using scRepertoire’s built-in TCR prediction functionality. This included grouping TCRs based on shared CDR3 motifs and sequence similarity to identify clonotypes potentially recognizing common antigens.

### Cell type identification and signatures

Single-cell transcriptomic and surface protein data were analyzed using Seurat (v5.0.1)^75^ to characterize major cell populations, including epithelial, stromal, B, T, and myeloid cells. UMAP embeddings were generated using Seurat’s Weighted Nearest Neighbor (WNN) workflow, enabling joint clustering based on both RNA and ADT modalities. For immune cells, dimensionality reduction and clustering were performed using WNN-based integration, whereas non-immune cells were analyzed using RNA-only PCA followed by UMAP. Clustering was carried out using the Louvain algorithm, applied to either the WNN graph (immune cells) or the RNA-derived Shared Nearest Neighbor (SNN) graph (non-immune cells). Clusters were initially assigned to broad compartments using canonical marker expression: epithelial cells (*EPCAM*, *KRT8*, *KRT18*), stromal cells (*CDH5*, *COL1A1*, *COL1A2*, *COL6A2*, *VWF*), T cells (*PTPRC*, *CD3D*, *CD3E*, *CD52*), B cells (*CD19*, *CD79A*, *CD79B*, *CD22*, *CD20*), and myeloid cells (*CD14*, *ITGAM*, *CD68*, *CSF1R*, *FCER1G*). Following compartment-level annotation, each population was subsetted and re-clustered to resolve finer cellular states. Cell type annotation was performed using a combination of automated label transfer via the SingleR package with curated reference datasets^78^, and manual refinement based on established literature and Human Cell Atlas markers^27–29,37,79^. Canonical marker gene expression profiles used for annotation are visualized in the corresponding figures.

### Differential cell-type abundance analysis using Milo

To assess differential cell-type abundance across experimental conditions, we employed the Milo R package^32^ on a WNN or SNN graph constructed from ExCITE-seq data. The precomputed graph from the Seurat object was converted into a Milo object using the buildFromAdjacency function, with parameters set to k = 20 and d = 25. Neighborhoods of transcriptionally similar cells were generated using the makeNhoods() function with graph-based refinement enabled (refined = TRUE, prop = 0.1, refinement_scheme = “graph”). The countCells function was then applied to quantify cell frequencies within each neighborhood per sample. To identify statistically significant changes in cell abundance, we performed differential abundance testing using testNhoods and graph-based multiple testing correction (fdr.weighting = “graph-overlap”). Neighborhoods with an adjusted spatial FDR < 0.1 were considered significantly differentially abundant and were visualized accordingly: red for decrease and blue for enrichment.

### Trajectory analysis

To model epithelial differentiation dynamics and predict lineage outcomes, we applied CellRank^31^, a probabilistic framework that reconstructs cellular trajectories by learning Markov chains from transcriptomic similarity structures. To infer pseudotime and differentiation potential, we used CytoTRACE^80^, which infers differentiation potential based on transcriptional entropy. The resulting pseudotime values were used to orient differentiation trajectories and identify candidate progenitor and terminal populations. A transition matrix representing cell-state progression probabilities was constructed using a k-nearest neighbor (kNN) graph derived from the transcriptomic space. Generalized Perron Cluster Cluster Analysis (GPCCA) was then used to decompose this transition matrix, enabling robust identification of initial and terminal states. Fate probabilities toward each terminal state were calculated for individual cells and visualized using UMAP embeddings and heatmaps. This enabled probabilistic lineage tracing and quantification of differentiation biases across epithelial subpopulations.

### Cell to cell communication and signaling activity

To infer intercellular signaling networks across defined cell populations, we utilized the LIANA (Ligand–Receptor Inference Analysis) framework, a consensus-based approach that integrates predictions from multiple established tools (e.g., CellPhoneDB, NATMI, and others)^35,81^. Putative ligand-receptor interactions were predicted using LIANA’s rank_aggregate function, which ranks and aggregates interaction scores across methods to identify statistically supported sender-receiver interactions based on ligand and receptor gene expression. To assess downstream pathway-level signaling, we integrated LIANA outputs with the decoupleR R package^82^. The run_ulm function was used to calculate pathway activity enrichment scores (e.g., for EGFR, WNT, TNFα, Hypoxia) by evaluating the expression of pathway-associated ligands in each cell type after subsetting by condition (CON, LS, and LS-CRC). Differential pathway activity across conditions was visualized using customized dot plots. Statistical significance was determined using Holm-adjusted p-values (p < 0.05).

To assess the differential expression of chemokines associated with MAIT cell trafficking, we analyzed expression levels of *CXCL12, CXCL16, CCL20, CXCL10,* and *CCL5* across epithelial subtypes using dot plot visualization. Expression patterns were evaluated to identify subset-specific transcriptional enrichment of these known MAIT-attracting chemokines, as previously described^38,53^

### Ro/e analysis

Ratio of observed to expected (Ro/e) quantifies whether a particular cell type is overrepresented or underrepresented in a given tissue or condition relative to what would be expected under a null hypothesis of uniform distribution. For each cell type, we first determined: Observed (Ro): The actual number of cells of a given subset detected in each condition or tissue; Expected (E): The number of cells expected under the assumption of proportional distribution across all groups, based on total cell counts. Mathematically: Ro/e = observed / expected^83^. An Ro/e > 1 indicates preferential enrichment of that cell subset in the tissue/condition, whereas Ro/e < 1 suggests relative depletion. A non-parametric pairwise Wilcoxon rank-sum test was employed to assess statistical significance of these deviations.

### Mice and AOM/DSS-induced CRC

All animal experiments were conducted according to the Public Health Service Policy on Humane Care and Use of Laboratory Animals, and were evaluated and approved by the Institutional Animal Care and Use Committee (IACUC) at the University of Pennsylvania (protocols #807356). We used 6 week old male C57BL/6J mice (Jackson Laboratory; stock number: 000664). Male mice were used because female mice display reduced tumor progression and lower mortality compared to males in the preclinical models of CRC^84,85^, precluding our ability to assess whether an intervention ameliorates tumorigenesis. Mice were treated with 5-OP-RU (20 µmol/L, gift from Dr. Oh, 10 mM stock in DMSO)^86^ with or without CpG (50 µg/mL, ODN 1585, InvivoGen), or vehicle control (PBS). Treatments were administered by concurrent oral gavage (200 µL) and intraperitoneal injection (200 µL) twice per week, beginning four weeks before and continuing throughout the duration of the AOM/DSS regimen. The azoxymethane (AOM)/dextran sodium sulfate (DSS) model was used to induce CRC, as previously described^87^. Briefly, the above mice received a single intraperitoneal injection of AOM (12 mg/kg in PBS, Sigma-Aldrich, Cat#A5486-25MG). Three days later, mice were administered 2.2% DSS (TdBLabs, Cat# 9011-18-1) in drinking water for five consecutive days, followed by 14 days of regular drinking water. This cycle was repeated three times. Cage exchange was performed every 3–5 days.

### Flow cytometry for patient samples

Single cell suspensions were prepared from human colonic mucosal biopsies as described above. Cells were washed in FACS buffer consisting of PBS supplemented with 1% BSA and 2 mM EDTA. Fc receptors were blocked using human TruStain FcX™ for 10 minutes on ice to minimize nonspecific antibody binding. Cells were then stained with fluorochrome-conjugated monoclonal antibodies against surface markers including EpCam, CD44, CD166, CD3, CD4, CD8, CD45, 5-OP-RU loaded MR1 tetramer, and other lineage-specific markers (see Table S9 for full antibody panel). Staining was carried out in the dark at 4°C for 30 minutes. Following staining, cells were washed three times with FACS buffer and filtered through a 70 μm cell strainer to remove aggregates. Samples were acquired on a spectral flow cytometer Aurora CS-1 and analyzed using FlowJo software (v11). Fluorescence-minus-one (FMO) controls were included to guide gating strategies. Data analysis was restricted to live and singlet populations.

### Single cells preparation for mouse colonic tissues

#### Tumor tissue

Tumor regions were carefully dissected and cut into small fragments (∼0.4 cm). Samples were incubated in 1 mL digestion buffer composed of RPMI 1640 supplemented with 10% FBS, 1% penicillin/streptomycin, 10 mM HEPES, collagenase type I (0.5 mg/mL), and DNase I (20 U/mL) for 40 minutes at 37°C. Following enzymatic digestion, tissues were mechanically dissociated using a syringe plunger, passed through a 100 µm cell strainer, and quenched with 15 mL cold complete RPMI (cRPMI). The resulting cell suspension was centrifuged at 400 g for 5 minutes and resuspended in FACS buffer for downstream applications.

#### Tumor-free colon tissue

Non-tumor colon segments (∼1 cm length) were initially incubated in 5 mL dissociation buffer composed of calcium-and magnesium-free PBS supplemented with 1 mM DTT (Sigma, Cat# D0632-5G) for 30 minutes at 37°C with shaking (200 rpm) to remove mucus. After washing thoroughly with PBS, tissues were minced into ∼0.5 cm pieces and digested in 5 mL digestion buffer (composition as above) for 30 minutes at 37°C with shaking (200 rpm). Digested suspensions were quenched with 15 mL cold cRPMI and filtered through a 100 µm strainer. The resulting single-cell suspension was centrifuged at 400 g for 5 minutes. To enrich immune cells, the pellet was resuspended in 40% Percoll (Sigma, Cat#P1644) and layered beneath 80% Percoll. Samples were centrifuged at 1,000 g for 22 minutes at room temperature without brake. After centrifugation, the top debris layer (∼3 mL) and bottom red blood cell-rich fraction (∼200 µL) were carefully removed. The remaining interphase layer was collected, washed with cRPMI, centrifuged at 700 g for 5 minutes, and resuspended in FACS buffer for flow cytometric analysis.

### Flow cytometry for mouse samples

Single cell suspensions prepared from mouse colon tissues (as described above) were washed and resuspended in FACS buffer. To minimize nonspecific antibody binding, Fc receptors were blocked using mouse TruStain FcX™ (BioLegend, Cat# 101319) for 10 minutes on ice. Cells were then stained with fluorochrome-conjugated monoclonal antibodies against surface markers, including EpCAM, CD3, CD4, CD8, CD45, TCRβ, MR1 tetramer, and other lineage-specific markers. The complete antibody panel is listed in Table S9. Surface staining was performed in the dark at 4°C for 30 minutes. Following incubation, cells were washed three times with FACS buffer and passed through a 70 μm cell strainer to remove clumps. Data acquisition was conducted using a spectral flow cytometer (Aurora CS-1, Cytek) and analyzed using FlowJo software (v11). Gating strategies were guided by fluorescence-minus-one (FMO) controls. Data analysis was restricted to live and singlet populations.

### Immunofluorescence staining and quantitative image analysis

Formalin-fixed, paraffin-embedded (FFPE) human colonic mucosal tissue sections were processed for immunofluorescence staining. After deparaffinization and rehydration, antigen retrieval was performed in sodium citrate buffer (Sigma, Cat#C9999) at 95°C for 30 minutes. Sections were cooled and blocked with 5% FBS and 0.1% Triton X-100(Sigma, Cat#T8787) in PBS for 1 hour at room temperature. Primary antibodies were incubated overnight at 4°C, targeting CCL20 (1:200, R&D Systems, Cat#AF360) and OLFM4 (1:150, Cell Signaling Technology, Cat# 14369T). After PBS washes, slides were incubated with secondary antibodies (anti-goat Alexa Fluor 800, 1:200, ThermoFisher, Cat# A32930; anti-rabbit Alexa Fluor 568, 1:200, ThermoFisher, Cat# A-11011) for 1 hour at room temperature. After PBS washes, slides were incubated with antibody against EpCAM (1:200, Abcam, Cat#ab237395) to label epithelial cells. Nuclei were counterstained with DAPI (1:2000, Bio-Rad), and sections were mounted using ProLong Gold Antifade (ThermoFisher, Cat#P36934). Imaging was conducted using a Leica confocal microscope at constant acquisition settings across all samples. Image quantification was performed in ImageJ (v1.54f). Individual fluorescence channels were analyzed separately. After manually selecting the DAPI positive area, binary thresholding was applied using standard intensity cutoffs (0–255), and the following parameters were extracted per channel: percent of positive area, mean intensity. CCL20 expression was normalized to the OLFM4+ signal by integrating intensity metrics and percent of positive area, enabling quantification of chemokine enrichment within epithelial stem-like compartments. At least 3-5 fields per sample were analyzed.

### Quantification and statistical analysis

All statistical analyses were conducted using base R and GraphPad Prism v9. For non-parametric data, group comparisons were performed using Dunn’s multiple comparisons test with Benjamini– Hochberg correction. For parametric datasets: Two-way ANOVA was used when datasets involved two independent variables, followed by Holm-Šidák post hoc tests; one-way ANOVA was used for comparisons involving a single independent variable, followed by Fisher’s LSD post hoc test. Correlation analyses were performed using simple linear regression. The significance of the regression slope was assessed using F-tests, and corresponding p-values were calculated in Prism v9. A threshold of p < 0.05 was considered statistically significant. Non-significant results (p > 0.05) were interpreted as a lack of meaningful linear association. All p-values are reported in the figure legends with the following notation: p < 0.05 (*), p < 0.01 (**), p < 0.001 (***), and “ns” for non-significant comparisons.

## Supporting information

None

## ACKNOWLEDGMENTS

We are deeply grateful to all the patients who generously donated biospecimens, making this study possible. We also thank Drs. John Wherry, Clifton Barry III, Manolis Roulis, Alexander C. Huang, and Divij Mathew for their valuable guidance and support. We also thank the CHOP Flow Cytometry Core, CHOP Single Cell Core, and CHOP Sequencing Core for assistance with experiments and the NIH Tetramer Core Facility (contract number 75N93020D00005) for providing tetramers. Additionally, we acknowledge the following University of Pennsylvania core facilities for their expert technical support: the Cell and Developmental Biology (CDB) Microscopy Core and the Molecular Pathology & Imaging Core (MPIC) (RRID: SCR_022420) supported by the UPENN Digestive and Liver Center (P30DK050306). This work was supported in part by the National Institutes of Health (NIH) grants DK093668 (K.C.), AI121244 (K.C.), AI140754 (K.C.), AI179896 (K.C.), and DK050306 (K.C.); pilot grant from King Center for Lynch Syndrome (K.C.), pilot grant from Abramson Cancer Center (K.C., B.K.), the Jason and Julie Borrelli Lynch Syndrome Research Fund (B.K.), and the King Family Fund for Lynch Syndrome Education, Outreach and Impact (B.K.). J.A. is supported by the Crohn’s and Colitis Foundation (#878246) the NIH NIDDK Diseases K23DK124570.

## AUTHOR CONTRIBUTIONS

H.Y., B.K., and K.C. conceived the study and interpreted the data. H.Y. and K.C. designed the experiments. B.K., M.D., J.A., M.D., B.C., R.K., and K.B. procured samples and information from human subjects. H.Y. performed all the experiments and bioinformatics data analyses. X.W., M.B., C.J.L, and N.B. assisted mouse experiments. S.O. generated 5-OP-RU. D.S. and S.B.K. assisted with computational analyses. H.Y., B.K., and K.C. wrote and revised the manuscript, with all authors contributing to its review and editing.

## DECLARATION OF INTERESTS

B.K. has received clinical research funding paid to the institution from Janssen, Immunovia, Freenome, Guardant, Epigenomics, Universal Diagnostics, and Recursion. B.K. is an advisory board member at Immunovia. K.C. has received research funding from Pfizer, Takeda, Pacific Biosciences, Genentech and Abbvie. K.C. has consulted for or received an honorarium for speaking engagement from Puretech Health, Genentech, Moderna, Gentibio, and Abbvie. K.C. is an inventor on US patent 10,722,600 and provisional patents 62/935,035 and 63/157,225. J.A. has received research grants from BioFire Diagnostics, Genentech, Janssen, and Takeda; consultancy fees, honorarium, or advisory board fees from Abbvie, Abivax, Adiso, Biomerieux, Bristol-Myers Squibb, Celltrion, Ferring, Fresenius, Janssen, Merck, Pfizer, Sanofi, Takeda, and Vedanta.

