## Supplementary material for "MAIT cell enrichment in Lynch syndrome is associated with immune surveillance and colorectal cancer risk": None

**SUPPLEMENTAL FIGURES AND LEGENDS**
**Figure S1**

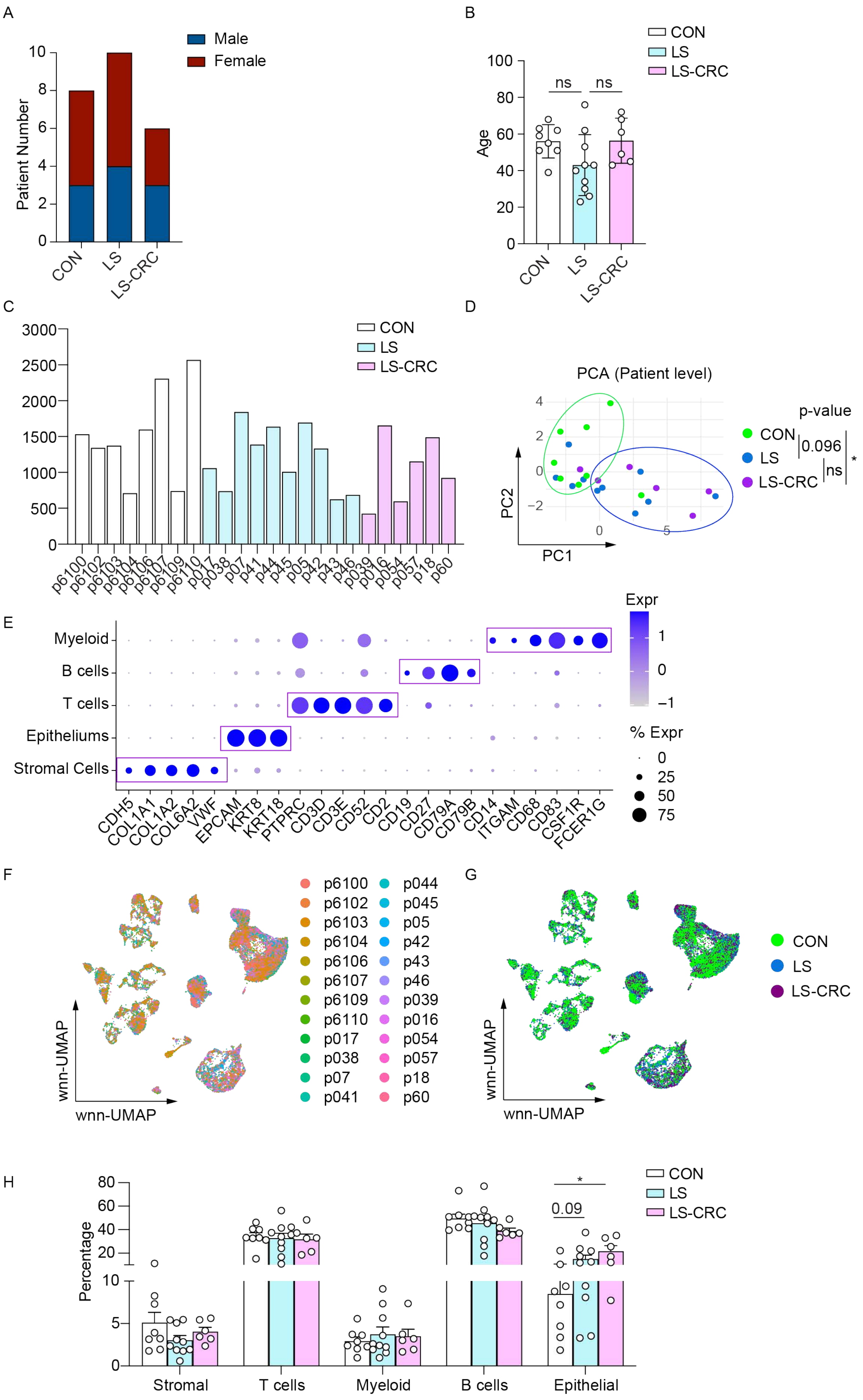

**Figure S1. Demographics of subjects and cellular landscape of human colorectal tissues revealed by ExCITE-seq profiling.**
(A) Distribution of male and female subjects across the three study groups: CON, LS, and LS-CRC (see Table S1).
(B) Comparison of individual age among the three groups (see Table S1).
(C) Total number of cells recovered per individual from ExCITE-seq, grouped by condition.
(D) Principal component analysis (PCA) plot showing individual-level clustering based on global transcriptional profiles. Each dot represents an individual and is colored by group: CON (green), LS (navy), and LS-CRC (purple). Statistical significance was assessed by PERMANOVA
(E) Dot plot displaying the expression of selected canonical marker genes across cell types. Dot size reflects the percentage of cells expressing each gene; color intensity indicates average log-normalized expression.
(F-G) UMAP projection of all cells colored by individual donors (F) or groups (G) (CON, LS, and LS-CRC.) shows broad overlap across conditions, indicating effective batch correction and integration across datasets.
(H) Proportions of five major cell compartments (stromal, T cells, myeloid, B cells, and epithelial cells) across CON, LS, and LS-CRC. Percentages are calculated relative to total cells per sample.

Dots represent one individual and bars show mean ± SEM for (B) and (H). Statistical significance was determined by two-way ANOVA (Holm-Šidák post hoc test, $*p < 0.05$, $**p < 0.001$, ns = not significant).

### Figure S2

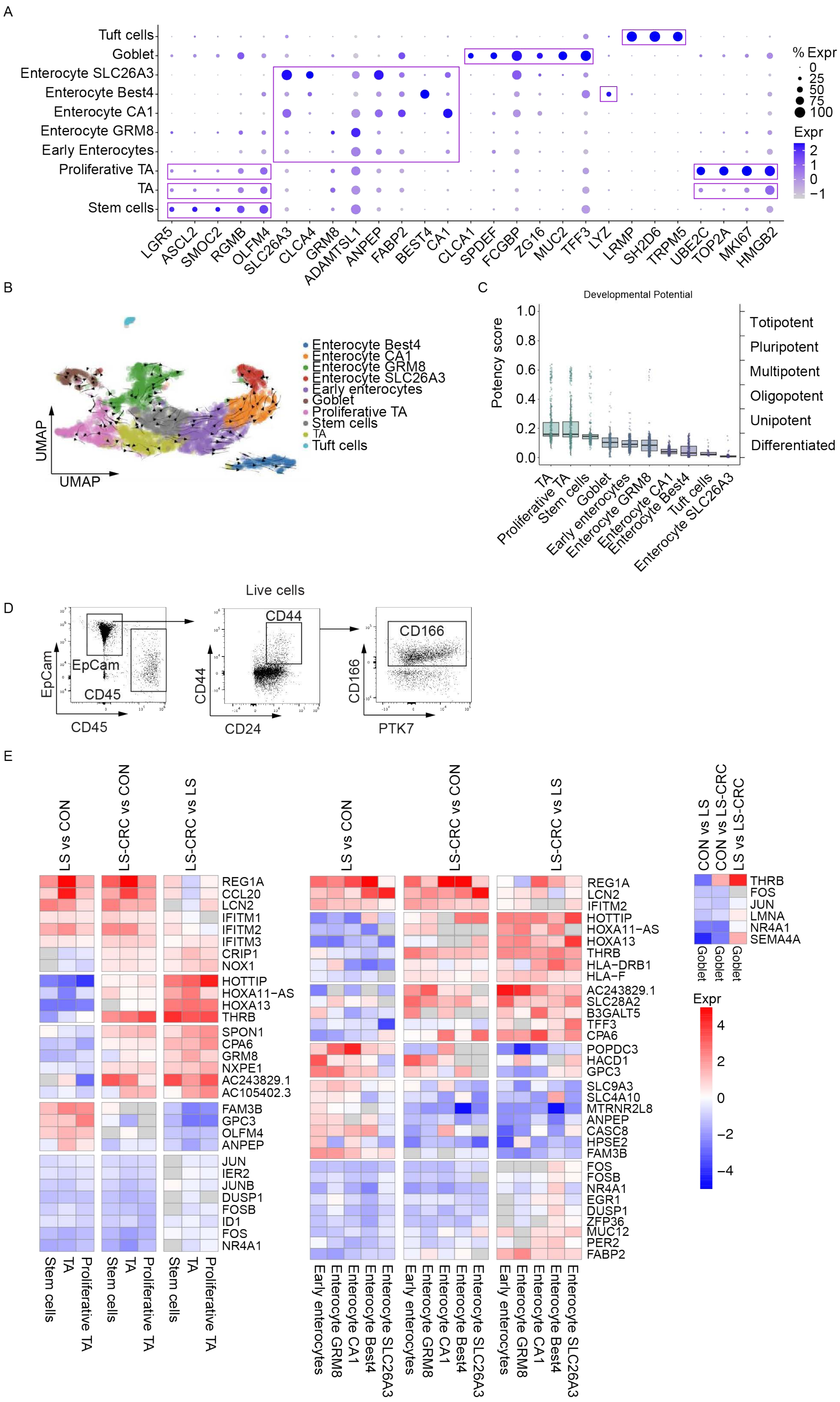

A
Tuft cells
Goblet
Enterocyte SLC26A3
Enterocyte Best4
Enterocyte CA1
Enterocyte GRM8
Early Enterocytes
Proliferative TA
TA
Stem cells
LGR5 ASCL2 SMOC2 RGMB OLFM4 SLC26A3 CLCA4 GRM8 ADAMTSL1 ANPEP FABP2 BEST4 CA1 CLCA1 SPDEF FCGBP ZG16 MUC2 TFF3 LYZ LRMP SH2D6 TRPM5 UBE2C TOP2A MKI67 HMGB2
% Expr
0
25
50
75
100
Expr
2
1
0
−1
B
Enterocyte Best4
Enterocyte CA1
Enterocyte GRM8
Enterocyte SLC26A3
Early enterocytes
Goblet
Proliferative TA
Stem cells
TA
Tuft cells
UMAP
UMAP
C
Developmental Potential
Potency score
1.0
0.8
0.6
0.4
0.2
0.0
Totipotent
Pluripotent
Multipotent
Oligopotent
Unipotent
Differentiated
TA
Proliferative TA
Stem cells
Goblet
Early enterocytes
Enterocyte GRM8
Enterocyte CA1
Enterocyte Best4
Tuft cells
Enterocyte SLC26A3
D
Live cells
EpCam
CD45
EpCam
CD45
CD44
CD44
CD24
CD166
CD166
PTK7
E
LS vs CON
LS-CRC vs CON
LS-CRC vs LS
REG1A
CCL20
LCN2
IFITM1
IFITM2
IFITM3
CRIP1
NOX1
HOTTIP
HOXA11−AS
HOXA13
THRB
SPON1
CPA6
GRM8
NXPE1
AC243829.1
AC105402.3
FAM3B
GPC3
OLFM4
ANPEP
JUN
IER2
JUNB
DUSP1
FOSB
ID1
FOS
NR4A1
Stem cells
TA
Proliferative TA
LS vs CON
LS-CRC vs CON
LS-CRC vs LS
REG1A
LCN2
IFITM2
HOTTIP
HOXA11−AS
HOXA13
THRB
HLA−DRB1
HLA−F
AC243829.1
SLC28A2
B3GALT5
TFF3
CPA6
POPDC3
HACD1
GPC3
SLC9A3
SLC4A10
MTRNR2L8
ANPEP
CASC8
HPSE2
FAM3B
FOS
FOSB
NR4A1
EGR1
DUSP1
ZFP36
MUC12
PER2
FABP2
Early enterocytes
Enterocyte GRM8
Enterocyte CA1
Enterocyte Best4
Enterocyte SLC26A3
CON vs LS
CON vs LS-CRC
LS vs LS-CRC
THRB
FOS
JUN
LMNA
NR4A1
SEMA4A
Goblet
Expr
4
2
0
−2
−4

**Figure S2. ExCITE-seq and flow cytometric analysis of epithelial cells in colorectal tissues.**
(A) Dot plot showing expression of canonical markers among indicated subtypes used to define epithelial cell types. Dot size indicates the percentage of cells expressing each gene; color intensity reflects average log-normalized expression.
(B) Pseudotime and fate probability map generated using CellRank, illustrating inferred trajectories from progenitor populations toward differentiated enterocyte subtypes.
(C) Developmental potential scores across epithelial subtypes, calculated using CytoTRACE.
(D) Flow cytometry gating strategies used to identify EpCAM+ cells and EpCAM+CD44+CD166+ cells representing epithelial cells and epithelial stem and progenitor cells, respectively.
(E) Top differentially expressed genes (DEGs) per epithelial subtype are shown for three comparisons: LS vs CON, LS-CRC vs CON, and LS-CRC vs LS. Rows represent genes, and columns represent epithelial subtypes. Color intensity denotes logFC values, with upregulation in red and downregulation in blue. Non-significant genes (p-value > 0.05) are shaded in gray. Full statistical results including adjusted p-values (Benjamini-Hochberg correction) are provided in Table S2.

### Figure S3

A

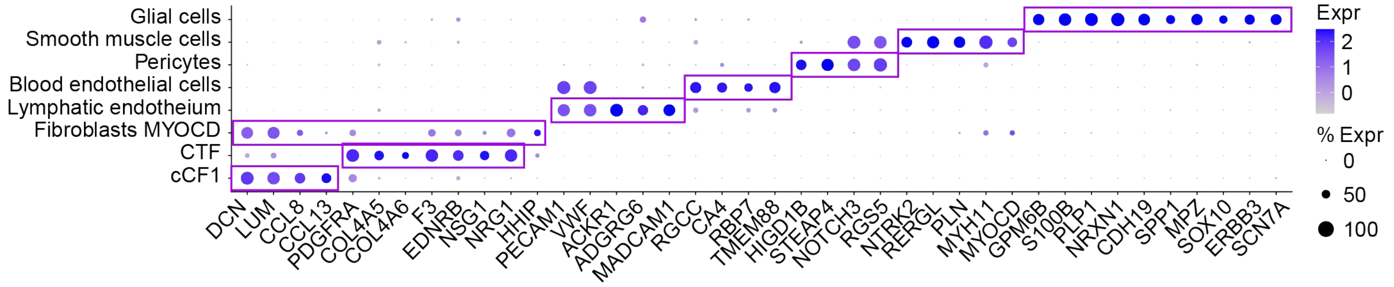


B

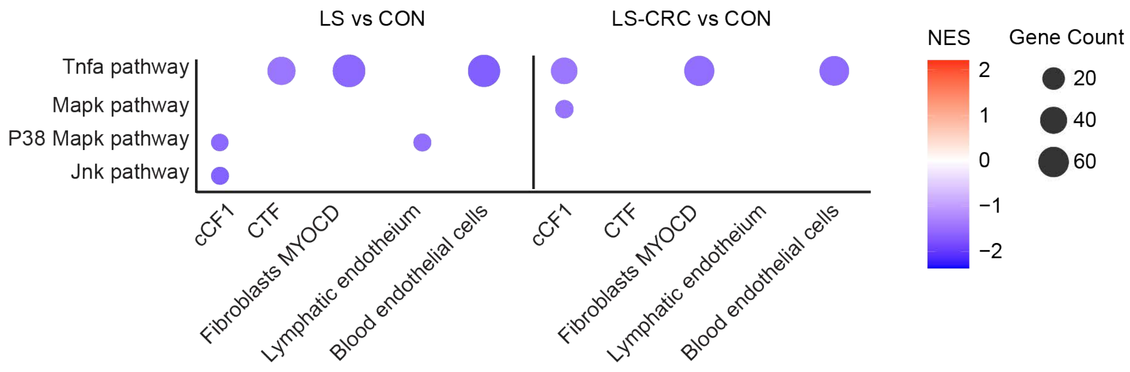


C

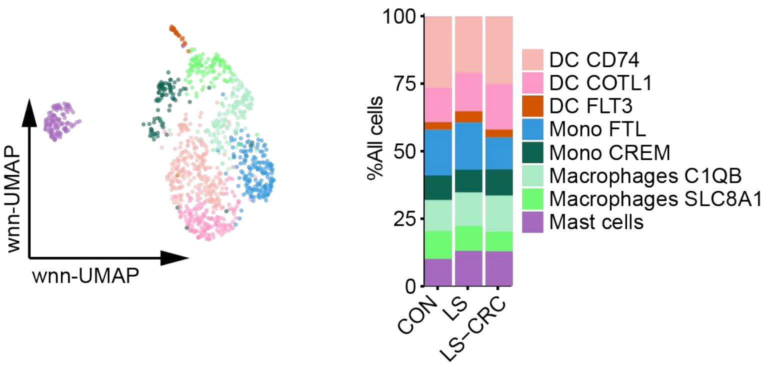


D

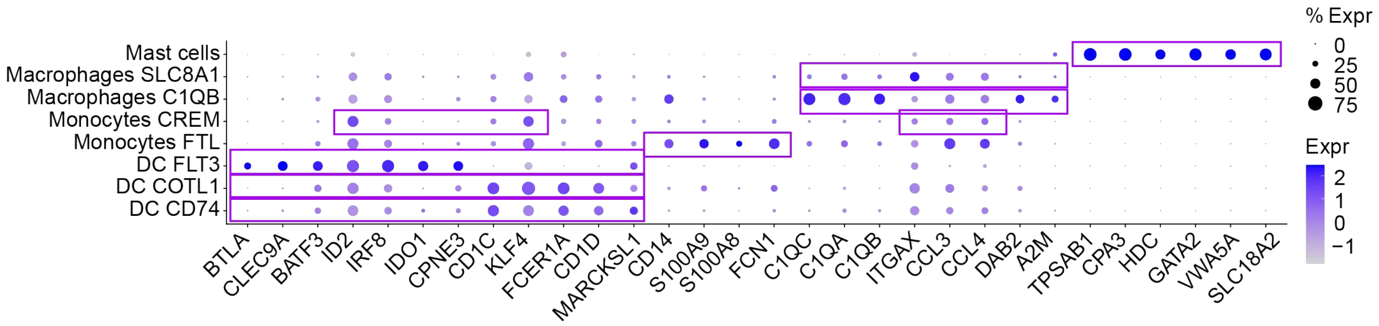

**Figure S3. Characterization of stromal and myeloid subtypes.**
(A) Dot plots showing canonical marker gene expression used to define stromal cell subtypes. Dot size represents the percentage of expressing cells, and color intensity reflects average log-normalized expression.
(B) GSEA of stromal cells comparing CON vs LS and CON vs LS-CRC. Each dot represents a differentially regulated pathway for a given stromal cell subtype, with color indicating the normalized enrichment score (NES) and dot size reflecting the number of genes contributing to the differential regulation (Gene count).
(C) wnn-UMAP projection (left) and bar graph showing relative abundance (right) of myeloid cells in CON, LS, and LS-CRC.
(D) Dot plots showing canonical marker gene expression used to define myeloid cell subtypes. Dot size represents the percentage of cells expressing the gene, and color intensity reflects average log-normalized expression.

**Figure S4**

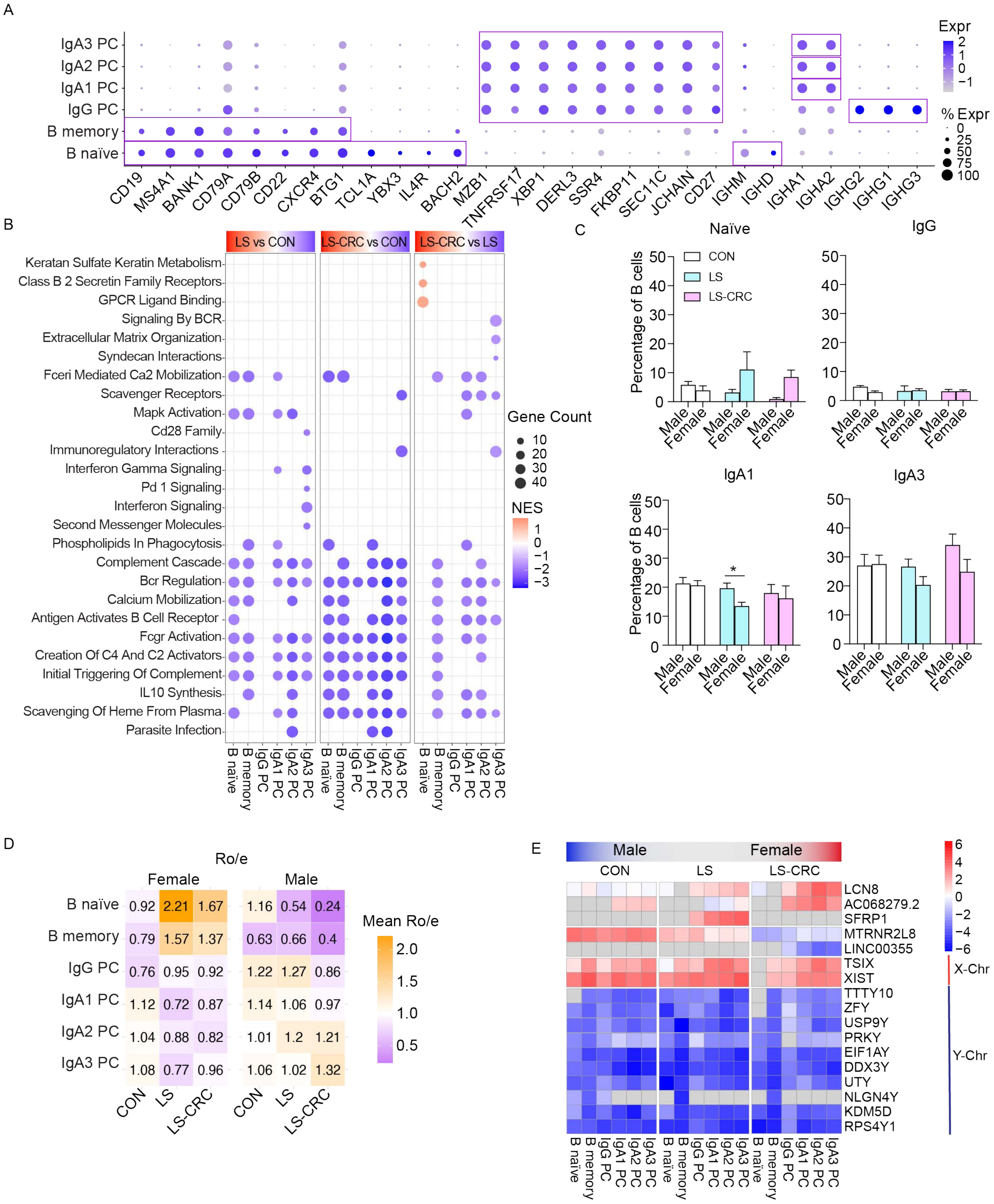
A
IgA3 PC
IgA2 PC
IgA1 PC
IgG PC
B memory
B naïve
CD19 MS4A1 BANK1 CD79A CD79B CD22 CXCR4 BTG1 TCL1A YBX3 IL4R BACH2 MZB1 TNFRSF17 XBP1 DERL3 SSR4 FKBP11 SEC11C JCHAIN CD27 IGHM IGHD IGHA1 IGHA2 IGHG2 IGHG1 IGHG3
Expr
% Expr
B
LS vs CON
LS-CRC vs CON
LS-CRC vs LS
Keratan Sulfate Keratin Metabolism
Class B 2 Secretin Family Receptors
GPCR Ligand Binding
Signaling By BCR
Extracellular Matrix Organization
Syndecan Interactions
Fceri Mediated Ca2 Mobilization
Scavenger Receptors
Mapk Activation
Cd28 Family
Immunoregulatory Interactions
Interferon Gamma Signaling
Pd 1 Signaling
Interferon Signaling
Second Messenger Molecules
Phospholipids In Phagocytosis
Complement Cascade
Bcr Regulation
Calcium Mobilization
Antigen Activates B Cell Receptor
Fcgr Activation
Creation Of C4 And C2 Activators
Initial Triggering Of Complement
IL10 Synthesis
Scavenging Of Heme From Plasma
Parasite Infection
Gene Count
NES
C
Naïve
IgG
IgA1
IgA3
CON
LS
LS-CRC
Percentage of B cells
Male
Female
D
Ro/e
Female
Male
Mean Ro/e
CON LS LS-CRC
E
Male
Female
CON
LS
LS-CRC
LCN8
AC068279.2
SFRP1
MTRNR2L8
LINC00355
TSIX
XIST
TTTY10
ZFY
USP9Y
PRKY
EIF1AY
DDX3Y
UTY
NLGN4Y
KDM5D
RPS4Y1
X-Chr
Y-Chr

**Figure S4. Characterization of B cells stratified by groups and sex.**

(A) Dot plots showing canonical marker gene expression used to define B cell subtypes. Dot size represents the percentage of cells expressing the gene and color intensity reflects average log-normalized expression.

(B) GSEA of B cells comparing CON vs LS and CON vs LS-CRC. Each dot represents an enriched pathway for a given B cell subtype, with color indicating NES and dot size reflecting the gene count. Enrichment was performed using curated gene sets related to immune and signaling pathways.

(C) Frequency of indicated B cell subtypes stratified by sex comparing CON, LS, and LS-CRC. Bars represent mean ± SEM. Statistical significance was determined by two-way ANOVA (Holm-Šidák post hoc test, $^{*}p < 0.05$, $^{**}p < 0.001$).

(D) Ratio of observed to expected cell number (Ro/e) scores for B cell subtypes stratified by sex. Values >1 indicate overrepresentation relative to expected frequencies. Scores are color-scaled by mean Ro/e per group.

(E) Top DEGs per B cell subtype stratified by sex comparing CON, LS, and LS-CRC. Rows represent genes, and columns represent B cell subtypes. Color intensity denotes logFC values, with upregulation in red and downregulation in blue. Non-significant genes ( p-value > 0.05) are shaded in gray. Full statistical results including adjusted p-values (Benjamini-Hochberg correction) are provided in Table S5.

### Figure S5

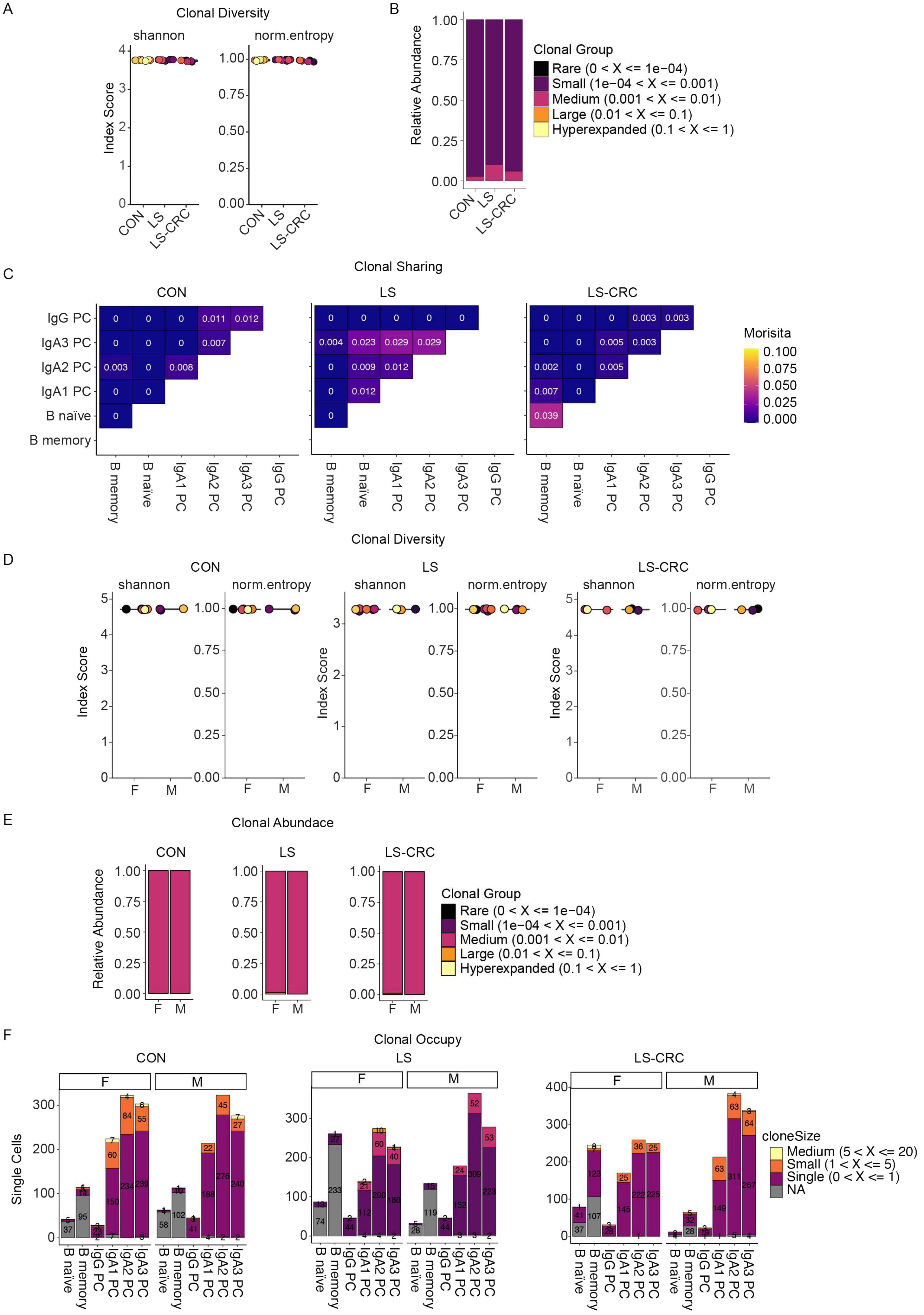
A
Clonal Diversity
shannon
norm.entropy
Index Score
CON
LS
LS-CRC
B
Relative Abundance
Clonal Group
Rare (0 < X <= 1e−04)
Small (1e−04 < X <= 0.001)
Medium (0.001 < X <= 0.01)
Large (0.01 < X <= 0.1)
Hyperexpanded (0.1 < X <= 1)
C
Clonal Sharing
Morisita
IgG PC
IgA3 PC
IgA2 PC
IgA1 PC
B naïve
B memory
D
Clonal Diversity
F
M
E
Clonal Abundace
F
Clonal Occupy
Single Cells
cloneSize
Medium (5 < X <= 20)
Small (1 < X <= 5)
Single (0 < X <= 1)
NA

**Figure S5. Characterization of B cell receptor (BCR) stratified by groups and sex.**
(A) Clonal diversity of BCR repertoires measured by Shannon entropy and normalized entropy (norm.entropy) in CON, LS, and LS-CRC groups. Values were calculated after downsampling to equal cell numbers across conditions (2,477 cells per group).
(B) Stacked bar plot showing the relative abundance of BCR clonotypes stratified by clonal size categories: Rare ($0 < X \leq 1e-4$), Small ($1e-4 < X \leq 0.001$), Medium ($0.001 < X \leq 0.01$), Large ($0.01 < X \leq 0.1$), and Hyperexpanded ($0.1 < X \leq 1$).
(C) Clonal sharing matrices based on the Morisita index comparing B cell subtypes within each group. Each heatmap displays the extent of clonal overlap between B cell subtypes (rows vs. columns), with color intensity and numeric values reflecting the degree of overlap.
(D) Clonal diversity of BCR repertoires measured by Shannon entropy and normalized entropy (norm.entropy) in CON, LS, and LS-CRC groups, stratified by sex. Diversity metrics were calculated after downsampling to control for sampling bias.
(E) Stacked bar plots showing the relative abundance of BCR clonotypes classified by clonal size categories and sex. Clonotypes are grouped into five categories based on their relative frequencies: Rare ($0 < X \leq 1e–4$), Small ($1e–4 < X \leq 0.001$), Medium ($0.001 < X \leq 0.01$), Large ($0.01 < X \leq 0.1$), and Hyperexpanded ($0.1 < X \leq 1$).
(F) Clonal occupancy of B cell subtypes in male and female individuals from each group. Bar plots show the number of single cells per clone size (NA, Single, Small, Medium) within each subset.

### Figure S6

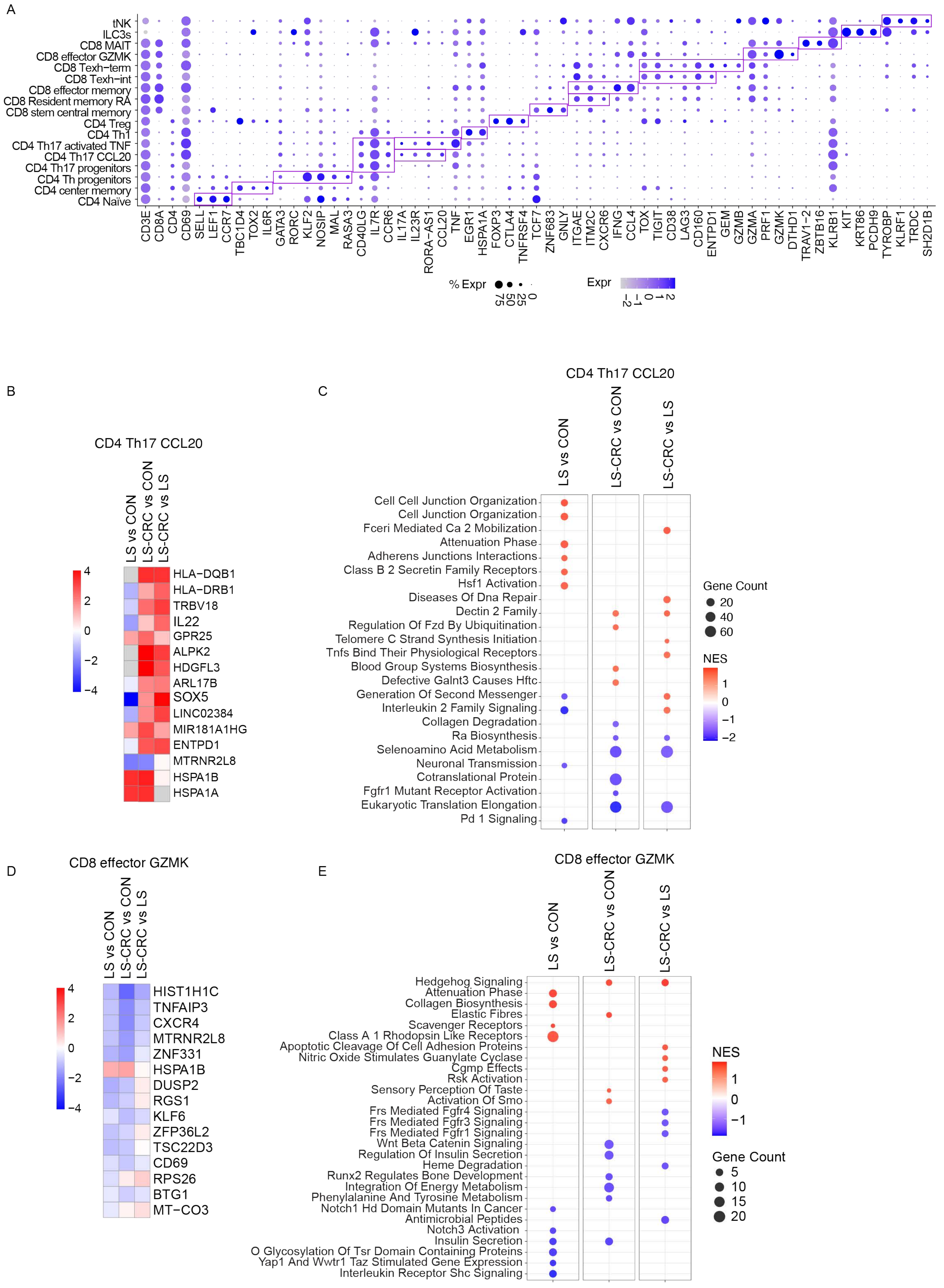
A
tNK
ILC3s
CD8 MAIT
CD8 effector GZMK
CD8 Texh−term
CD8 Texh−int
CD8 effector memory
CD8 Resident memory RA
CD8 stem central memory
CD4 Treg
CD4 Th1
CD4 Th17 activated TNF
CD4 Th17 CCL20
CD4 Th17 progenitors
CD4 Th progenitors
CD4 center memory
CD4 Naïve
CD3E CD8A CD4 CD69 SELL LEF1 CCR7 TBC1D4 TOX2 IL6R GATA3 RORC KLF2 NOSIP MAL RASA3 CD40LG IL7R CCR6 IL17A IL23R RORA−AS1 CCL20 TNF EGR1 HSPA1A FOXP3 CTLA4 TNFRSF4 TCF7 ZNF683 GNLY ITGAE ITM2C CXCR6 IFNG CCL4 TOX TIGIT CD38 LAG3 CD160 ENTPD1 GEM GZMB GZMA PRF1 GZMK DTHD1 TRAV1−2 ZBTB16 KLRB1 KIT KRT86 PCDH9 TYROBP KLRF1 TRDC SH2D1B
% Expr 0 25 50 75
Expr 2 1 0 −1 −2
B
CD4 Th17 CCL20
LS vs CON
LS-CRC vs CON
LS-CRC vs LS
4 2 0 −2 −4
HLA−DQB1
HLA−DRB1
TRBV18
IL22
GPR25
ALPK2
HDGFL3
ARL17B
SOX5
LINC02384
MIR181A1HG
ENTPD1
MTRNR2L8
HSPA1B
HSPA1A
C
CD4 Th17 CCL20
LS vs CON
LS-CRC vs CON
LS-CRC vs LS
Cell Cell Junction Organization
Cell Junction Organization
Fceri Mediated Ca 2 Mobilization
Attenuation Phase
Adherens Junctions Interactions
Class B 2 Secretin Family Receptors
Hsf1 Activation
Diseases Of Dna Repair
Dectin 2 Family
Regulation Of Fzd By Ubiquitination
Telomere C Strand Synthesis Initiation
Tnfs Bind Their Physiological Receptors
Blood Group Systems Biosynthesis
Defective Galnt3 Causes Hftc
Generation Of Second Messenger
Interleukin 2 Family Signaling
Collagen Degradation
Ra Biosynthesis
Selenoamino Acid Metabolism
Neuronal Transmission
Cotranslational Protein
Fgfr1 Mutant Receptor Activation
Eukaryotic Translation Elongation
Pd 1 Signaling
Gene Count
20
40
60
NES
1
0
−1
−2
D
CD8 effector GZMK
LS vs CON
LS-CRC vs CON
LS-CRC vs LS
4 2 0 −2 −4
HIST1H1C
TNFAIP3
CXCR4
MTRNR2L8
ZNF331
HSPA1B
DUSP2
RGS1
KLF6
ZFP36L2
TSC22D3
CD69
RPS26
BTG1
MT−CO3
E
CD8 effector GZMK
LS vs CON
LS-CRC vs CON
LS-CRC vs LS
Hedgehog Signaling
Attenuation Phase
Collagen Biosynthesis
Elastic Fibres
Scavenger Receptors
Class A 1 Rhodopsin Like Receptors
Apoptotic Cleavage Of Cell Adhesion Proteins
Nitric Oxide Stimulates Guanylate Cyclase
Cgmp Effects
Rsk Activation
Sensory Perception Of Taste
Activation Of Smo
Frs Mediated Fgfr4 Signaling
Frs Mediated Fgfr3 Signaling
Frs Mediated Fgfr1 Signaling
Wnt Beta Catenin Signaling
Regulation Of Insulin Secretion
Heme Degradation
Runx2 Regulates Bone Development
Integration Of Energy Metabolism
Phenylalanine And Tyrosine Metabolism
Notch1 Hd Domain Mutants In Cancer
Antimicrobial Peptides
Notch3 Activation
Insulin Secretion
O Glycosylation Of Tsr Domain Containing Proteins
Yap1 And Wwtr1 Taz Stimulated Gene Expression
Interleukin Receptor Shc Signaling
NES
1
0
−1
Gene Count
5
10
15
20

**Figure S6. Characterization and transcriptional profiling of T cell subtypes.**
(A) Dot plot displaying expression of canonical marker genes used to define T cell subtypes. Dot size represents the percentage of expressing cells, and color intensity reflects average log-normalized expression.
(B,D) Heatmaps showing the DEGs in CD4 Th17 CCL20 (B) and CD8 effector GZMK (D) cells comparing LS vs CON, LS-CRC vs CON, and LS-CRC vs LS. Color indicates logFC, with red indicating upregulation and blue indicating downregulation. Non-significant changes ($p > 0.05$) are shown in gray. Full statistical results including adjusted p-values (Benjamini-Hochberg correction) are provided in Table S6.
(C,E) GSEA for CD4 Th17 CCL20 (C) and CD8 effector GZMK (E) cells comparing LS vs CON, LS-CRC vs CON, and LS-CRC vs LS. Each dot represents an enriched signaling pathway, with color denoting the NES and size corresponding to the gene count in the pathway.

**Figure S7**

A

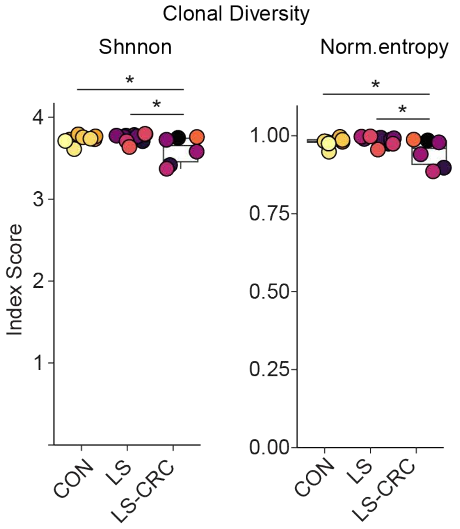


C

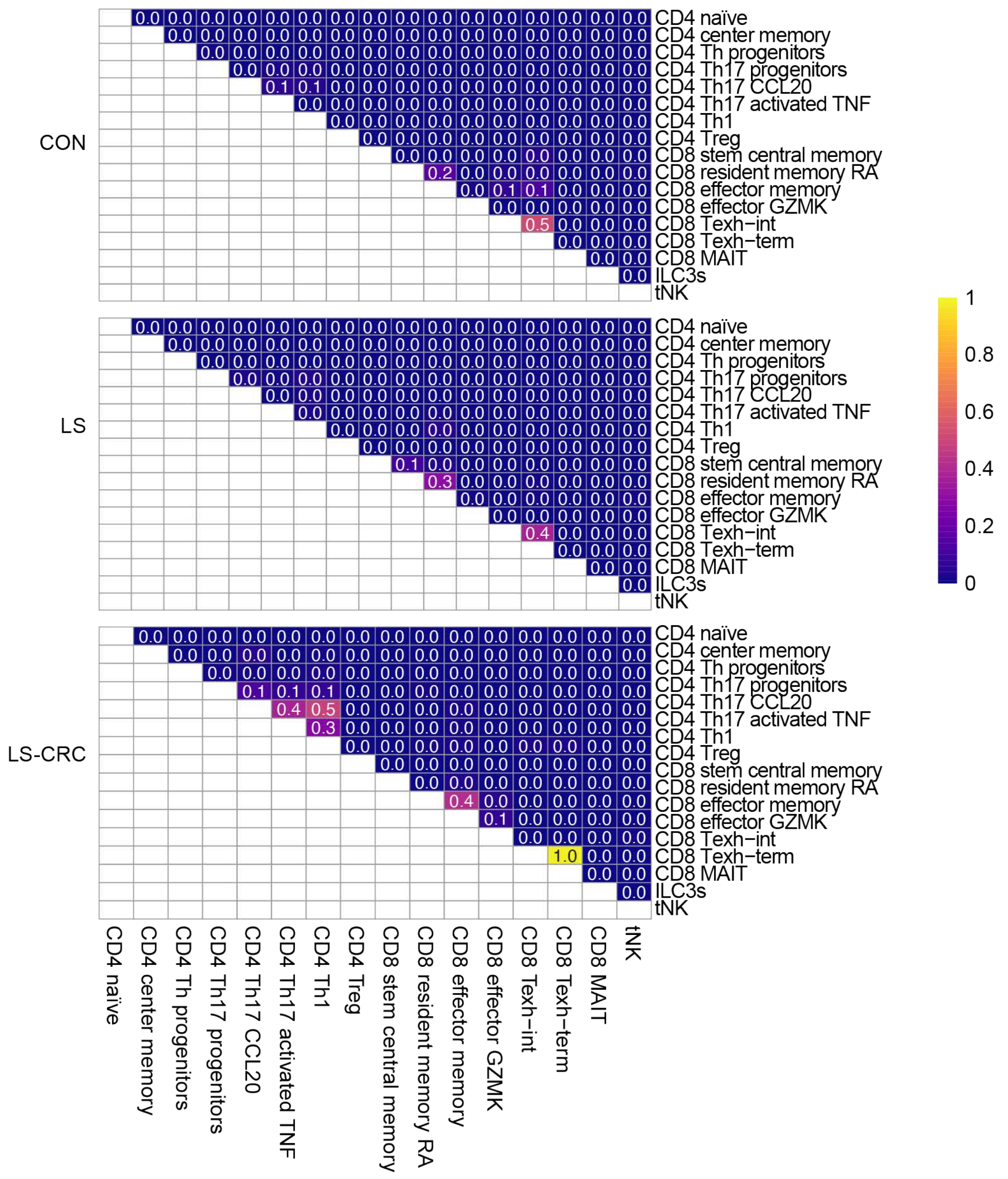


B

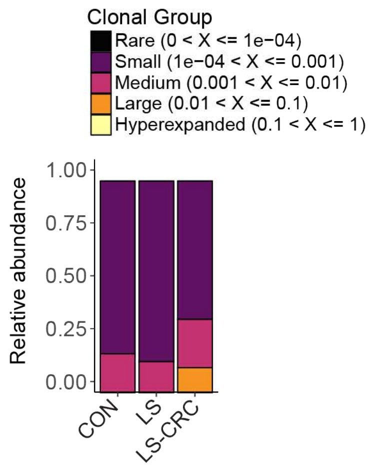


D

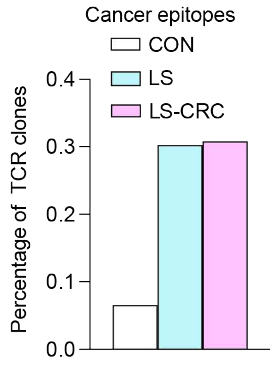

**Figure S7. Characterization of T cell receptor (TCR) repertoire and clonal sharing across groups.**
(A) Clonal diversity of TCR repertoires in CON, LS, and LS-CRC groups, assessed by Shannon entropy and normalized entropy (norm.entropy). All values were computed following downsampling to equal T cell numbers per group (2,191 cells per group). Statistical comparisons between groups were performed using Dunn's multiple comparisons test (Benjamini-Hochberg correction, (*$p < 0.05$)
(B) Stacked bar plot showing the relative abundance of TCR clonotypes stratified by clonal size categories: Rare ($0 < X \leq 1e-4$), Small ($1e-4 < X \leq 0.001$), Medium ($0.001 < X \leq 0.01$), Large ($0.01 < X \leq 0.1$), and Hyperexpanded ($0.1 < X \leq 1$).
(C) Clonal sharing matrices based on Morisita index values comparing TCR repertoire overlap between each T cell subtype (rows vs. columns) within CON, LS, and LS-CRC groups. Color intensity and numeric annotations reflect the extent of repertoire overlap.
(D) Proportion of T cells with TCR complementarity determining region 3 (CDR3) predicted to recognize tumor-associated antigens in the three study groups using Epitope Database (ScRepertoire).

**Figure S8**

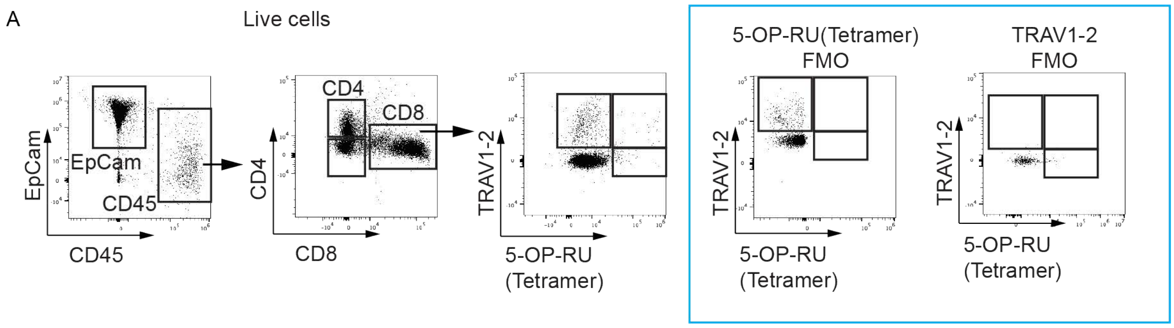


**Figure S8. Flow cytometry gating strategy for identification of MAIT cells**
(A) Representative flow cytometry plots depicting sequential gating steps from live cells.

**Figure S9**

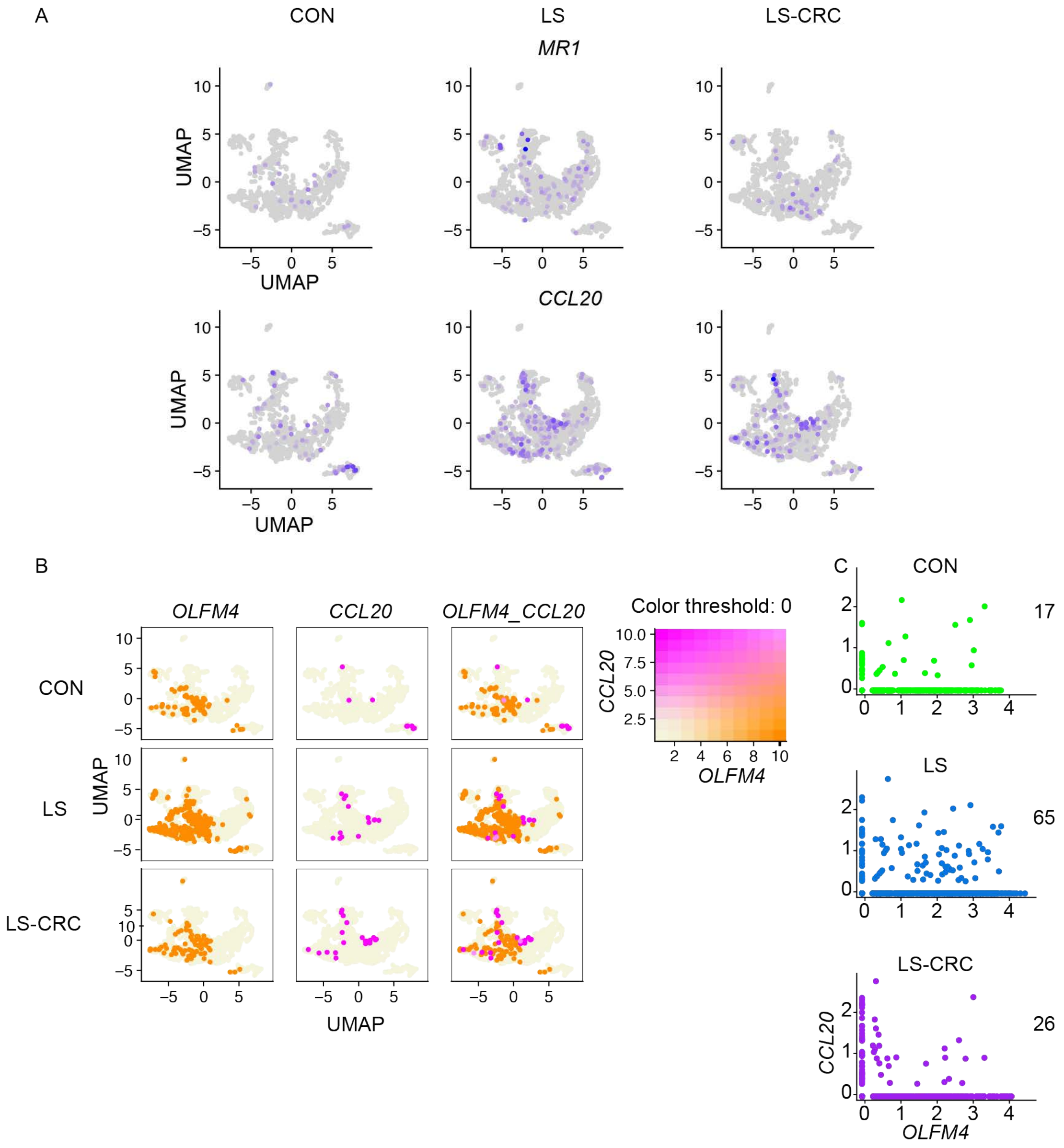


**Figure S9. Co-expression of CCL20 and OLFM4 identifies epithelial progenitor subsets in Lynch syndrome tissues**

(A) UMAP overlays showing expression of MR1 (top row) and CCL20 (bottom row) in epithelial cells from each group. Increased CCL20 expression is noted in LS and LS-CRC tissues.

(B) UMAP projections highlighting expression of OLFM4, CCL20, and their co-expression (OLFM4+ CCL20+) in epithelial cells across conditions. Co-expressing cells were colored based on scaled gene expression intensity. A bivariate color threshold plot (right) was used to define OLFM4+CCL20+ double-positive populations.

(C) Scatter plots showing single-cell co-expression of CCL20 and OLFM4 with counts of double-positive cells indicated (right) after downsampling (493 epithelial stem and progenitor cells per group). LS tissues show the highest frequency of OLFM4+CCL20+ epithelial cells, suggesting enrichment of inflammatory stem/progenitor states in pre-malignant lesions.

**Figure S10**

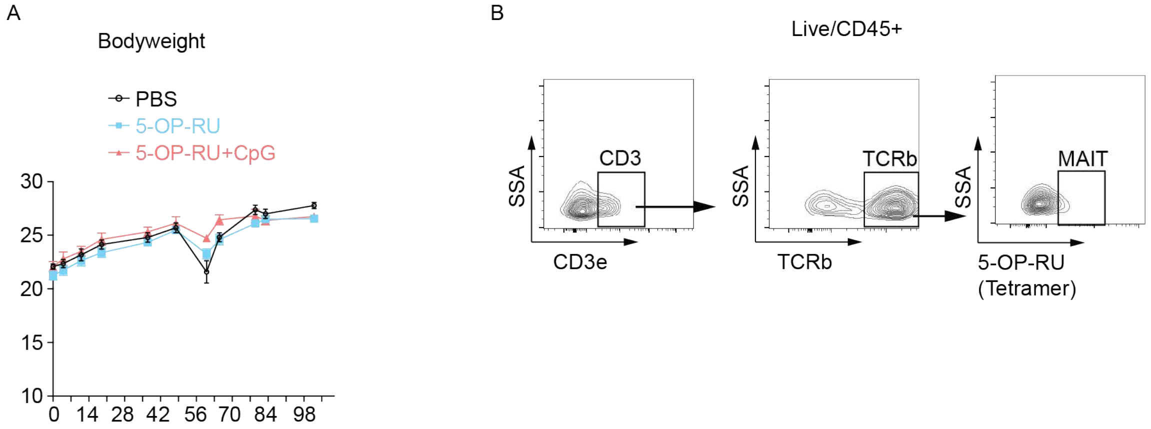


**Figure S10. Characterization of mice treated with 5-OP-RU and AOM/DSS**

(A) Longitudinal bodyweight measurements of mice treated with PBS, 5-OP-RU, or 5-OP-RU + CpG across the duration of the AOM/DSS experimental timeline. No significant differences in bodyweight were observed among treatment groups.

(B) Representative flow cytometry plots illustrating the sequential gating strategy used to identify MAIT cells. Gating was performed on live CD45+ cells, followed by selection of CD3+ T cells, TCRβ+ lymphocytes, and 5-OP-RU tetramer-positive (MAIT) cells.
